# Cold-inducible GOT1 activates the malate-aspartate shuttle in brown adipose tissue to support fuel preference for fatty acids

**DOI:** 10.1101/2024.11.18.623867

**Authors:** Chul-Hong Park, Minsung Park, Miranda E. Kelly, Helia Cheng, Sang Ryeul Lee, Cholsoon Jang, Ji Suk Chang

**Author notes:** Address correspondence to: Ji Suk Chang, Pennington Biomedical Research Center, 6400 Perkins Road, Baton Rouge, Louisiana 70808, USA.

## Abstract

Brown adipose tissue (BAT) simultaneously metabolizes fatty acids (FA) and glucose under cold stress but favors FA as the primary fuel for heat production. It remains unclear how BAT steer fuel preference toward FA over glucose. Here we show that the malate-aspartate shuttle (MAS) is activated by cold in BAT and plays a crucial role in promoting mitochondrial FA utilization. Mechanistically, cold stress selectively induces glutamic-oxaloacetic transaminase (GOT1), a key MAS enzyme, via the β-adrenergic receptor-PKA-PGC-1α axis. The increase in GOT1 activates MAS, transferring reducing equivalents from the cytosol to mitochondria. This process enhances FA oxidation in mitochondria while limiting glucose oxidation. In contrast, loss of MAS activity by GOT1 deficiency reduces FA oxidation, leading to increased glucose oxidation. Together, our work uncovers a unique regulatory mechanism and role for MAS in mitochondrial fuel selection and advances our understanding of how BAT maintains fuel preference for FA under cold conditions.

**Highlights:**

- *Got1* is markedly induced by cold in BAT via a β-adrenergic receptor-PKA-PGC-1α axis
- The increase in cytosolic GOT1 activates the malate-aspartate shuttle (MAS)
- MAS activation promotes fatty acid oxidation while reducing glucose oxidation
- Loss of MAS activity in BAT by *Got1* deletion shifts the fuel preference to glucose

## Introduction

Brown adipocytes, located in the interscapular brown adipose tissue (BAT), are a distinct type of fat cell that dissipates nutrient-derived energy in the form of heat via mitochondrial uncoupling protein 1 (UCP1) ^1–3^. Brown-like beige adipocytes also emerge within white adipose tissue during prolonged cold exposure or pharmacological stimulation of β_3_-adrenergic receptors (β_3_AR) ^4–10^. The presence of brown/beige adipocytes has been established in adult humans ^10–15^, and their activity is associated with increased energy expenditure and improved systemic lipid and glucose homeostasis. These properties make brown/beige adipocytes an appealing target against obesity and its related metabolic disorders.

BAT simultaneously metabolizes a variety of carbon substrates, such as fatty acids, glucose, and amino acids to meet thermogenic and biosynthetic needs under cold stress ^1,16^. Fatty acids (FA), released from intracellular triacylglycerol (TAG) stores or taken up from the circulation, are rapidly broken down into acetyl-CoA via FA β-oxidation with a concomitant production of NADH and FADH_2_ as byproducts. Acetyl-CoA enters the TCA cycle where it is further oxidized to generate NADH and FADH_2_. The resultant NADH and FADH_2_ supply electrons to the electron transport chain (ETC), creating a proton gradient across the inner mitochondrial membrane (IMM). UCP1 residing in the IMM allows these protons to flow back into the mitochondrial matrix, thus leading to heat production rather than ATP synthesis ^2,3^. As fatty acids (FA) produce more energy than any other substrates (e.g., 31 NADH and 15 FADH_2_ from one palmitic acid molecule compared to 6 NADH and 2 FADH_2_ from one glucose molecule), FA is the primary fuel source for UCP1-mediated thermogenesis. Moreover, FA directly activates UCP1 by inducing a conformational change that allows protons to be transported across the IMM ^17^. Cold-activated BAT also takes up a large amount of circulating glucose; however, glucose is only modestly (<15%) used as a fuel source for thermogenesis ^18–21^. Increasing evidence indicates that the majority of glucose-derived carbons produced in glycolysis are released as lactate ^20,21^ or channeled into various biosynthetic pathways, such as glycogen synthesis, pentose phosphate pathway, and glycerol-3-phosphate synthesis, which are essential for replenishment of intracellular TAG in lipid droplets ^20–23^. The glycolytic end product pyruvate entering the mitochondria is primarily used to replenish TCA cycle intermediates (i.e., oxaloacetate) via pyruvate carboxylation ^23^ or directed toward *de novo* fatty acid synthesis (FAS) ^21,24^ although a small pool of pyruvate enters the TCA cycle as acetyl-CoA. Branched-chain amino acids (BCAA) are used as nitrogen donors for biosynthesis of non-essential amino acids ^25^ and as additional, but minor, carbon sources for TCA cycle intermediates ^26^. This distinct substrate utilization likely allows BAT to efficiently meet the dual demands of heat production and cellular biosynthesis in response to cold stress. However, the regulatory mechanisms by which BAT prioritizes FA as the primary fuel for thermogenesis while directing other substrates to meet various metabolic needs still remain elusive.

The malate-aspartate shuttle (MAS) is an important biochemical system that maintains cytosolic and mitochondrial NAD^+^/NADH redox balance under glycolytic conditions. MAS transports reducing equivalents produced in glycolysis from the cytosol to mitochondria ^27^, since the mitochondrial inner membrane (IMM) is impermeable to NADH. In cells with high energy demands, MAS enables glycolysis-derived NADH to be fed into the ETC for ATP production, while also regenerating NAD^+^ in the cytosol to sustain glycolysis. MAS is involved in various tissue-specific metabolic processes, such as neurotransmitter synthesis in the brain ^28^, pumping activity of the heart ^29^, urea cycle in the liver ^30^, insulin secretion by pancreatic β-cells ^31^, and exercise capacity in skeletal muscle under aerobic conditions ^32^. However, MAS’s role in BAT with high metabolic and thermogenic demands remains unclear. MAS is composed of two mitochondrial carriers (OGC, AGC) and two pairs of cytosolic and mitochondrial enzymes – glutamic-oxaloacetic transaminases (cGOT1, mGOT2) and malate dehydrogenases (cMDH1, mMDH2). Intriguingly, we found that GOT1 is markedly induced by cold in BAT from its barely detectable level, while other MAS enzymes remain unchanged. Thus, we sought to investigate whether the increase in GOT1 protein levels activates MAS in BAT and what role MAS plays in BAT under cold conditions. By using BAT-specific *Got1* knock-in and *Got1* knockout mice and their respective brown adipocytes, we demonstrate that brown adipocyte GOT1 is a critical node linking cold-stimulated β-adrenergic signaling to the MAS, and cold-activated MAS is essential for promoting efficient utilization of FA over glucose in the mitochondria. This discovery expands our understanding of MAS beyond NADH transfer, positioning it as a critical regulator of mitochondrial fuel selection in BAT.

## Results

### GOT1, a key enzyme in the malate-aspartate shuttle, is highly elevated by cold in BAT

The malate-aspartate shuttle (MAS) utilizes two pairs of cytosolic and mitochondrial enzymes and two mitochondrial carriers (Fig. 1A). MAS is active in energy-demanding tissues, such as the brain, heart, liver, and to a lesser extent, skeletal muscle, and plays a critical role in cellular metabolism in a tissue-specific manner ^27^. Our transcriptome analysis ^33^ of BAT from mice housed at room temperature or exposed to 4°C for 6h unexpectedly revealed that the *Got1* gene, but not other MAS enzyme genes (*Got2*, *Mdh1*, *Mdh2*), is highly upregulated by cold (Fig. 1B). Likewise, mice stimulated with a cold-mimetic β_3_-adrenergic receptor agonist CL316243 ^5–7^ showed increased expression of *Got1* in BAT with no change in *Got2*, *Mdh1*, and *Mdh2* gene expression (Fig. 1C), suggesting that cold-stimulated βAR signaling promotes the transcription of the *Got1* gene. Although GOT1 was abundantly expressed in many tissues like the brain, liver, heart, skeletal muscle, and kidney, GOT1 protein was barely detectable in BAT at room temperature (Fig. 1D). In agreement with the gene expression data, cold exposure markedly increased GOT1 protein levels (Fig. 1E), which were also coupled with heightened enzymatic activity (Fig. 1F). However, the protein levels of other MAS enzymes were relatively unchanged during cold exposure (Fig. 1E). Similar to BAT, GOT1 protein was hardly detectable in inguinal white adipose tissue (IWAT) at room temperature (Fig. 1D and 1G), but we observed a significant induction of GOT1 in IWAT undergoing browning during prolonged cold exposure (Fig. 1G). Together, these data indicate that cold stress upregulates *Got1* gene expression in brown and beige adipocytes. In support of our findings, a recent brown, beige, and white adipocyte transcriptome analysis in mice and humans ^34^ identified *Got1* as a highly enriched gene in cold-activated brown and beige adipocytes.

**Figure 1.**
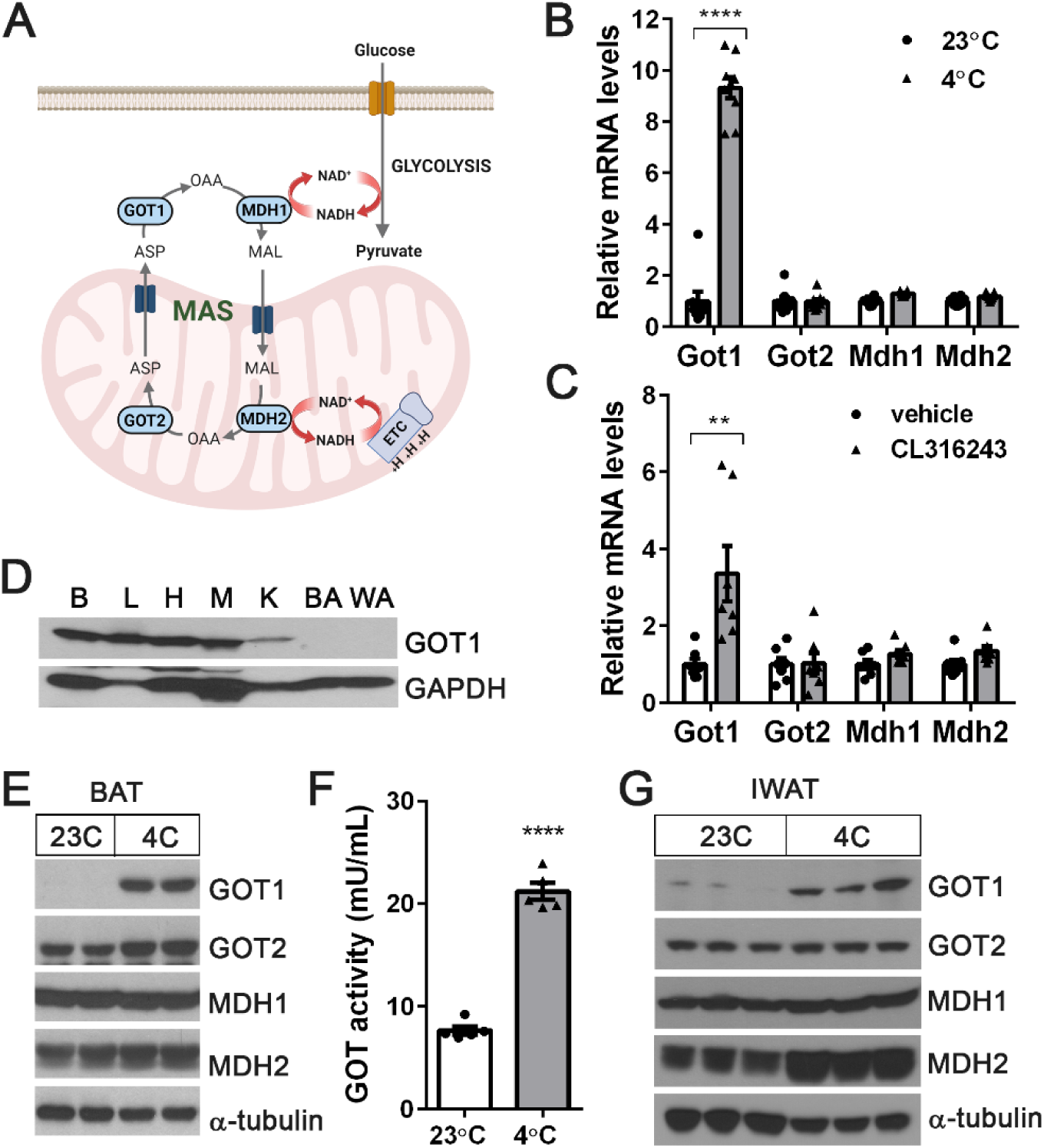
Cold-dependent induction of GOT1, a key enzyme of the malate-aspartate shuttle in BAT. (A) A schematic of the malate-aspartate shuttle (MAS), which consists of two pairs of cytosolic (GOT1 and MDH1) and mitochondrial (GOT2 and MDH2) enzymes and two mitochondrial transporters. MAS transports reducing equivalents produced in glycolysis from the cytosol to mitochondria in the form of malate, which is oxidized back to oxaloacetate (OAA), regenerating NADH in the mitochondria. (B) qPCR analysis to determine the effect of cold stress on MAS gene expression in BAT. BL6 mice were housed at 23°C or exposed to 4°C for 5h. (C) Effect of β_3_-adrenergic stimulation on MAS gene expression in BAT. BL6 mice were administered with vehicle or a β_3_AR agonist CL316243 (1 mg/kg BW) for 5h. (D) Expression of GOT1 protein in brain (B), liver (L), heart (H), muscle (M), kidney (K), brown adipose (BA), and white adipose (WA) tissue. (E) Cold-dependent induction of GOT1 protein in BAT. (F) Increased GOT activity in cold-activated BAT. (G) Effect of cold stress on the MAS enzyme levels in IWAT. BL6 mice were housed at 23°C or exposed to 4°C for 7 days. All data are presented as the Mean ± SEM. \*\**p*<0.01, \*\*\*\**p*<0.0001.

### The *Got1* gene expression is upregulated via a βAR-cAMP-PKA-PGC-1α/NT-PGC-1α axis

Cold stress stimulates the sympathetic nervous system to release norepinephrine, which binds to β-adrenergic receptors on brown adipocytes and activates a signaling cascade leading to increased expression of a number of genes including *Ucp1* ^1^. To verify that brown adipocyte β-adrenergic signaling is responsible for increased *Got1* gene expression in BAT, we stimulated fully differentiated brown adipocytes ^35^ with a βAR agonist isoproterenol or a PKA activator cAMP. Both significantly induced *Got1* gene expression with no effect on *Got2*, *Mdh1*, and *Mdh2* genes (Fig. 2A and 2B). Given that transcriptional co-activators, full-length PGC-1α and its shorter isoform NT-PGC-1α, regulate a number of cold-responsive genes in BAT in response to βAR signaling ^35,36^, we asked whether *Got1* is a downstream target of PGC-1α and/or NT-PGC-1α. Our BAT ChIP-seq ^33^ with PGC-1α antibody detecting both PGC-1α and NT-PGC-1α ^35,36^ revealed that PGC-1α/NT-PGC-1α are enriched at the *GOT1* gene promoter that contains two consensus DNA binding motifs for estrogen-related receptors (ERRs) (Fig. 2C). This is consistent with the previous finding that ERRα/γ binds to and activates the *GOT1* gene promoter in murine heart ^37^. Indeed, we were able to confirm the recruitment of PGC-1α/NT-PGC-1α to one of the ERRE sites at the *GOT1* gene promoter (−604/-596) (Fig. 2D). Given that PGC-1α and NT-PGC-1α are transcriptional co-activators of ERRs in brown adipocytes ^36,38,39^, the βAR-PKA-PGC-1α/NT-PGC-1α-ERRs axis likely promotes *Got1* transcription in brown adipocytes. While the loss of both PGC-1α and NT-PGC-1α in *Pgc-1α*^-/-^ brown adipocytes ^40,41^ blunted *Got1* gene expression, the deletion of either PGC-1α or NT-PGC-1α in brown adipocytes ^41,42^ did not alter its expression (Fig. 2E). Likewise, re-expression of PGC-1α or NT-PGC-1α in *Pgc-1α*^-/-^ brown adipocytes was sufficient to promote *Got1* gene expression (Fig. 2F). Together, these results delineate that cold stress upregulates *Got1* gene expression in brown adipocytes via the well-established βAR-cAMP-PKA-PGC-1α/NT-PGC-1α-ERR pathway (Fig. 2G).

**Figure 2.**
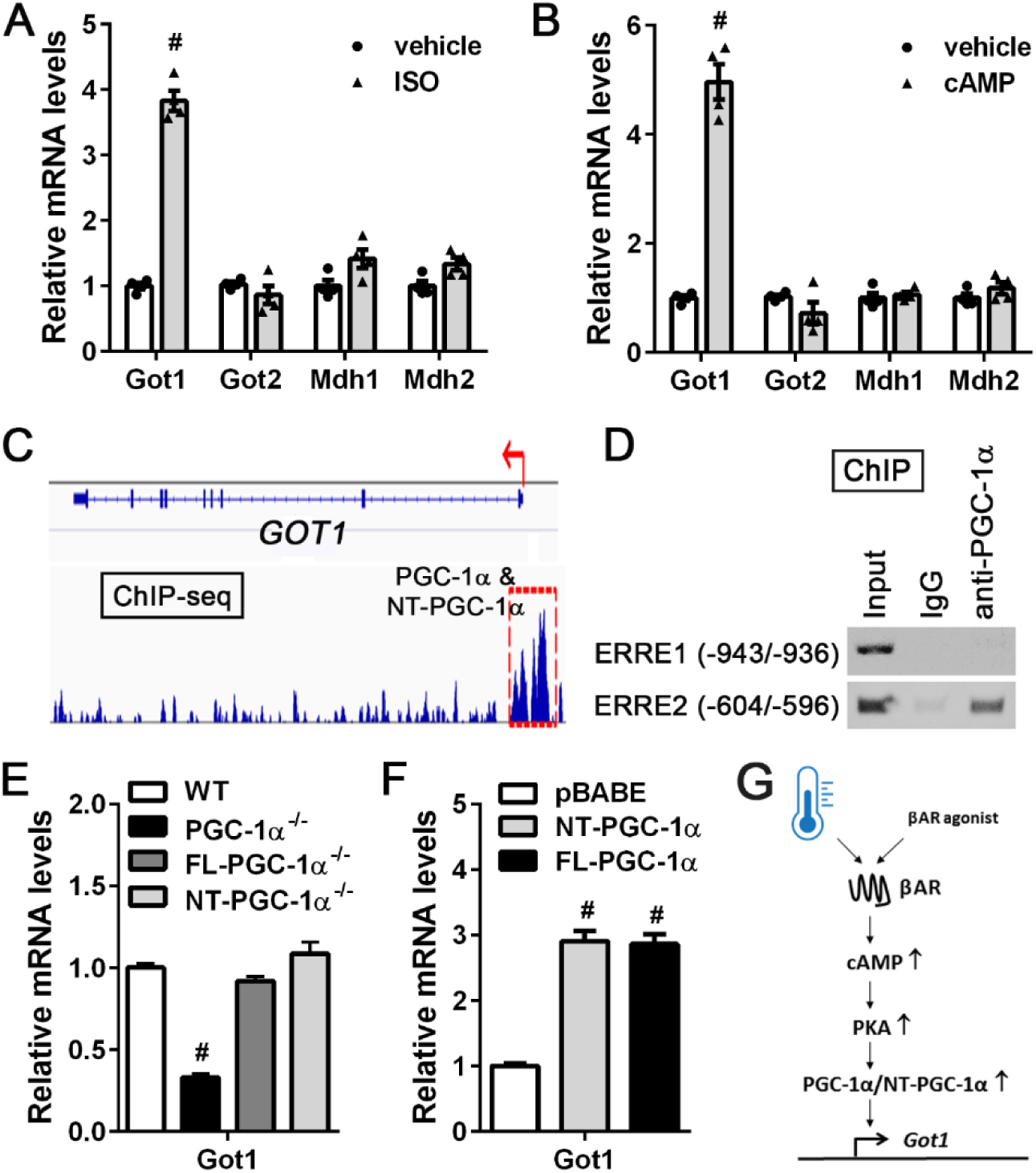
Cold-stimulated βAR signaling promotes the transcription of the *GOT1* gene through the βAR-cAMP-PKA-PGC-1α/NT-PGC-1α axis. (A, B) Upregulation of *Got1* gene expression in brown adipocytes treated with 5 µM isoproterenol (ISO) or 1 mM dibutyryl cAMP for 4h. (C) Enrichment of PGC-1α/NT-PGC-1α at the *GOT1* gene promoter in cold-activated BAT. The PGC-1α/NT-PGC-1α ChIP-seq peak ^33^ was visualized by the Integrative Genome Viewer (IGV) v2.3. (D) PCR analysis of PGC-1α/NT-PGC-1α recruitment to the ERRE of the *GOT1* promoter. ChIP was carried out with PGC-1α antibody in nuclear extracts of BAT extracted from mice exposed to 4°C for 5h. (E) Effect of single or combined loss of full-length (FL)-PGC-1α and NT-PGC-1α on *Got1* gene expression. Fully differentiated brown adipocytes were treated with 1 mM dibutyryl cAMP for 4h (n=4/group). (F) Effect of ectopic expression of FL-PGC-1α or NT-PGC-1α on *Got1* expression in brown adipocytes (n=4/group). (G) Schematic presentation of transcriptional regulation of the *Got1* gene by cold or β-AR agonist in brown adipocytes. All data are presented as the Mean ± SEM. ^#^*p*<0.0001.

### Mice overexpressing GOT1 in BAT exhibit enhanced BAT thermogenesis

This finding led us to ask whether GOT1 acts as a molecular switch for activating the MAS in BAT in response to cold. To address this question, we generated a *Got1* knock-in mouse line where the transgene of *CAG-loxP-neo-3xpA(stop)-loxP-Got1* was inserted into the Rosa26 locus (Fig. 3A). While the polyadenylation sequence terminates transcription initiated by the CAG promoter, the Cre-mediated excision of the stop cassette enables targeted expression of *Got1*. R26-*Got1*^fl/+:UCP1-cre/+^ (R26-*Got1*^BATOE/+^) mice were generated by crossing R26-*Got1*^fl/fl^ mice with *Ucp1*-Cre mice ^43^ that express Cre recombinase in BAT at room temperature. R26-*Got1*^BATOE/+^ mice housed at 23°C exhibited selective expression of *Got1* in BAT, but not in other tissues (Fig. 3B). Moreover, GOT1 protein levels produced by the R26-*Got1*^BATOE/+^ allele at 23°C were comparable to those induced by cold in R26-*Got1*^fl/+^ control mice (Fig. 3C), and GOT1 overexpression did not alter the expression of other MAS enzymes (Fig. 3D), thereby validating the overexpression of GOT1 in BAT without cold exposure. Brown adipocyte morphology was relatively normal in R26-*Got1*^BATOE/+^ mice (Fig. 3E), but GOT1 overexpression caused a significant change in the cellular NAD^+^/NADH ratio in BAT (Fig. 3F), as expected from GOT1’s role in MAS activation.

**Figure 3.**
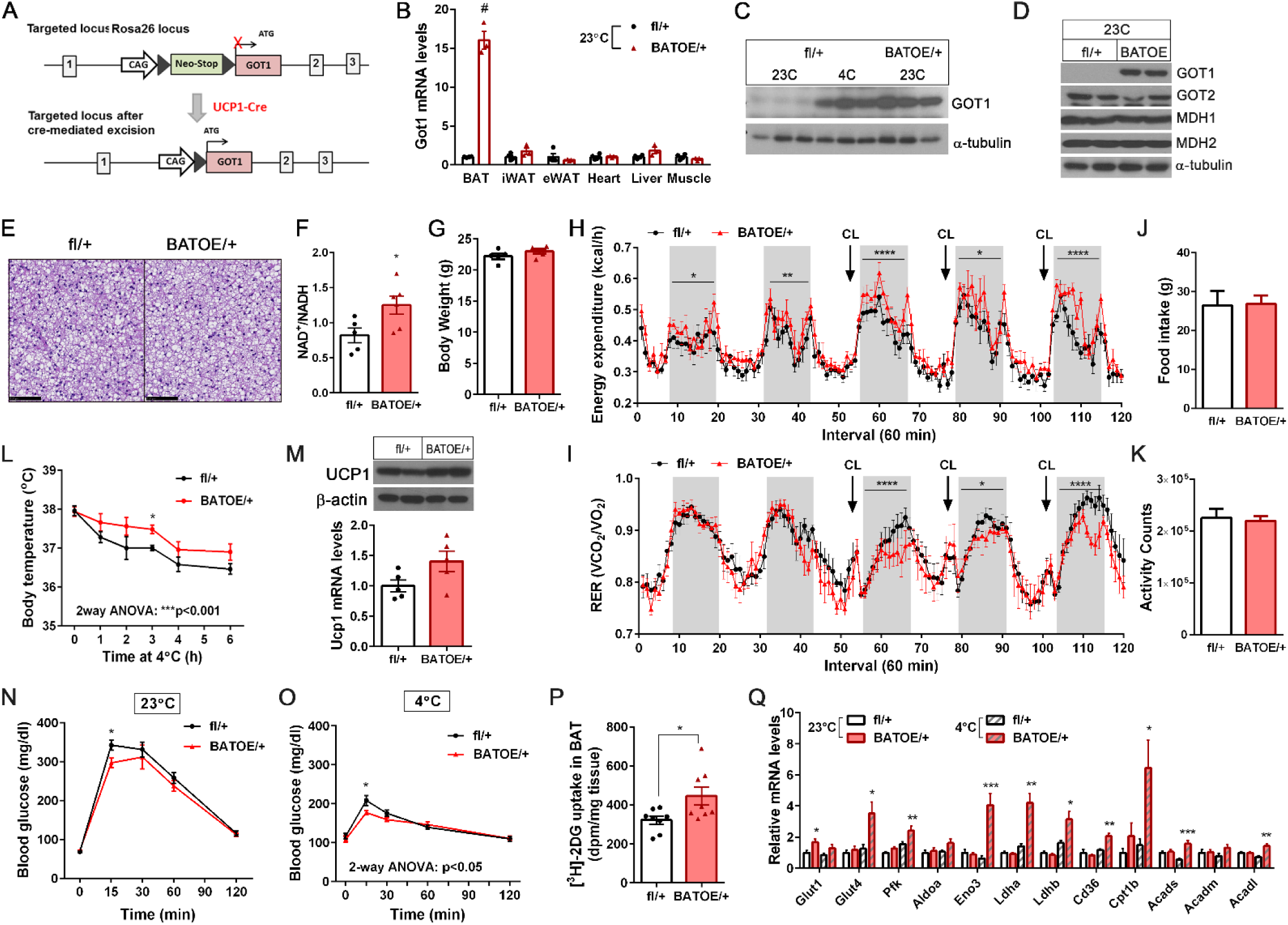
BAT-specific overexpression of GOT1 in mice enhances BAT thermogenesis. (A) Generation of BAT-specific *Got1* knock-in mice. The transgene of *CAG-loxP-neo-3xpA(stop)-loxP-Got1* was inserted into the Rosa26 locus. Cre-mediated excision of the stop cassette enables targeted expression of *Got1* in BAT. (B) qPCR analysis for validation of BAT-specific *Got1* expression in R26-*Got1*^BATOE/+^ mice housed at 23°C. (C) Cold-independent elevation of GOT1 protein in BAT of R26-*Got1*^BATOE/+^ male mice at 23°C. (D) No alteration of other MAS enzyme expression by GOT1 overexpression in BAT. (E) H&E staining of BAT from mice at 23°C. Black bars represent 100 µm. (F) Measurement of NAD^+^/NADH ratios in BAT homogenates. (G-K) Measurement of body weight, energy expenditure, RER, food intake, and locomotor activity in R26-*Got1*^fl/+^ and R26-*Got1*^BATOE/+^ female mice (n=5/group). Mice were placed in indirect calorimetric chambers and monitored prior to and during administration of a β_3_AR agonist CL316243 (1mg/kg BW, daily). (L) Improved cold tolerance of R26-*Got1*^BATOE/+^ female mice (n=5/group). The body temperature was measured with a rectal thermometer. (M) Ucp1 mRNA and protein levels in BAT. (N, O) Glucose tolerance tests of R26-*Got1*^fl/+^ and R26-*Got1*^BATOE/+^ female mice at 23°C (n=7/group) and at 4°C (n=7/group). (P) Measurement of [^3^H]-2DG uptake by BAT extracted from mice housed at 23°C. (Q) qPCR analysis of BAT extracted from R26-*Got1*^fl/+^ and R26-*Got1*^BATOE/+^ mice housed at 23°C or 4°C (n=6-7/group). All data are presented as the Mean ± SEM. \**p*<0.05, \*\**p*<0.01, \*\*\**p*<0.001, \*\*\*\**p*<0.0001, ^#^*p*<0.0001.

To investigate the impact of GOT1 overexpression on BAT thermogenesis, we evaluated whole-body energy expenditure (EE) by indirect calorimetry in R26-*Got1*^BATOE/+^ and R26-*Got1*^fl/+^ mice prior to and after stimulation of BAT with a selective β_3_AR agonist CL316243 ^5–7^. Body weights were not different between the groups prior to indirect calorimetry (Fig. 3G). R26-*Got1*^BATOE/+^ mice exhibited slightly higher basal EE compared to littermate controls and displayed a greater increase in EE immediately after administration of CL316243 (Fig. 3H).

Intriguingly, increased EE was associated with a lower respiratory exchange ratio (RER) (Fig. 3I), reflecting preferential use of fat as the energy source for BAT thermogenesis. Food intake and locomotor activity did not differ between the groups during indirect calorimetry (Fig. 3J and 3K). In line with higher EE, R26-*Got1*^BATOE/+^ mice exhibited improved cold tolerance upon exposure to 4°C (Fig. 3L). The cold tolerance phenotype was associated with increasing trends in UCP1 mRNA and protein levels in *Got1*OE BAT (Fig. 3M). In line with lower RER, *Got1*OE BAT showed elevated expression of genes linked to FA uptake (*Cd36*), mitochondrial FA transport (*Cpt1b*), and FA β-oxidation (*Acads*, *Acadl*) (Fig. 3Q). Together, these data suggest that an increased energy supply for thermogenesis through FA oxidation might account for enhanced BAT thermogenesis in R26-*Got1*^BATOE/+^ mice.

BAT is the tissue with highest rates of glucose uptake during cold exposure ^44–46^, contributing to enhanced glucose tolerance during GTT ^47^. Thus, we assessed glucose tolerance in R26-*Got1*^BATOE/+^ and R26-*Got1*^fl/+^ mice at room temperature and during cold exposure. At room temperature, a rise in blood glucose levels at 15 min was lower in R26-*Got1*^BATOE/+^ mice compared to littermate controls, although overall glycemic differences between the groups were not statistically significant (Fig. 3N). BAT extracted from R26-*Got1*^BATOE/+^ mice housed at room temperature indeed showed higher uptake of [^3^H]-2-deoxyglucose (2-DG) (Fig. 3P). During cold exposure, R26-*Got1*^BATOE/+^ mice exhibited improved glucose tolerance compared to littermate controls (Fig. 3O). In line with these data, *Got1*OE BAT showed augmented expression of genes involved in glucose uptake (*Glut1*, *Glut4*) and glycolysis (*Pfk*, *Eno3*, *Ldha/b*) (Fig. 3Q).

Collectively, these data show that GOT1 overexpression in BAT boosts BAT’s ability to adapt to cold by increasing fat oxidation and glucose uptake and metabolism.

### GOT1 overexpression in brown adipocytes activates the malate-aspartate shuttle (MAS)

Next, we sought to investigate whether the observed phenotypes of R26-*Got1*^BATOE/+^ mice indeed result from GOT1-mediated activation of MAS in brown adipocytes. Thus, we generated *Got1*OE brown adipocytes by isolating stromal vascular fraction (SVF) from R26-*Got1*^fl/fl^ BAT, followed by Cre-mediated recombination ^48^ and differentiation into brown adipocytes. GOT1 overexpression did not alter brown adipogenesis as evidenced by no difference in adipogenic gene expression (Fig. 4A). The increase in cytosolic GOT1 levels resulted in increased GOT1 enzyme activity in the cytosolic fraction (Fig. 4B and 4D), and cytosolic GOT1 activity was completely inhibited by aminooxyacetate (AOA) that inhibits GOT1 and GOT2 ^49^ (Fig. 4C and 4D). In the MAS, GOT1 catalyzes the conversion of aspartate to oxaloacetate, which is reduced to malate via malate dehydrogenase (MDH1) with the coupled oxidation of NADH to NAD^+^.

**Figure 4.**
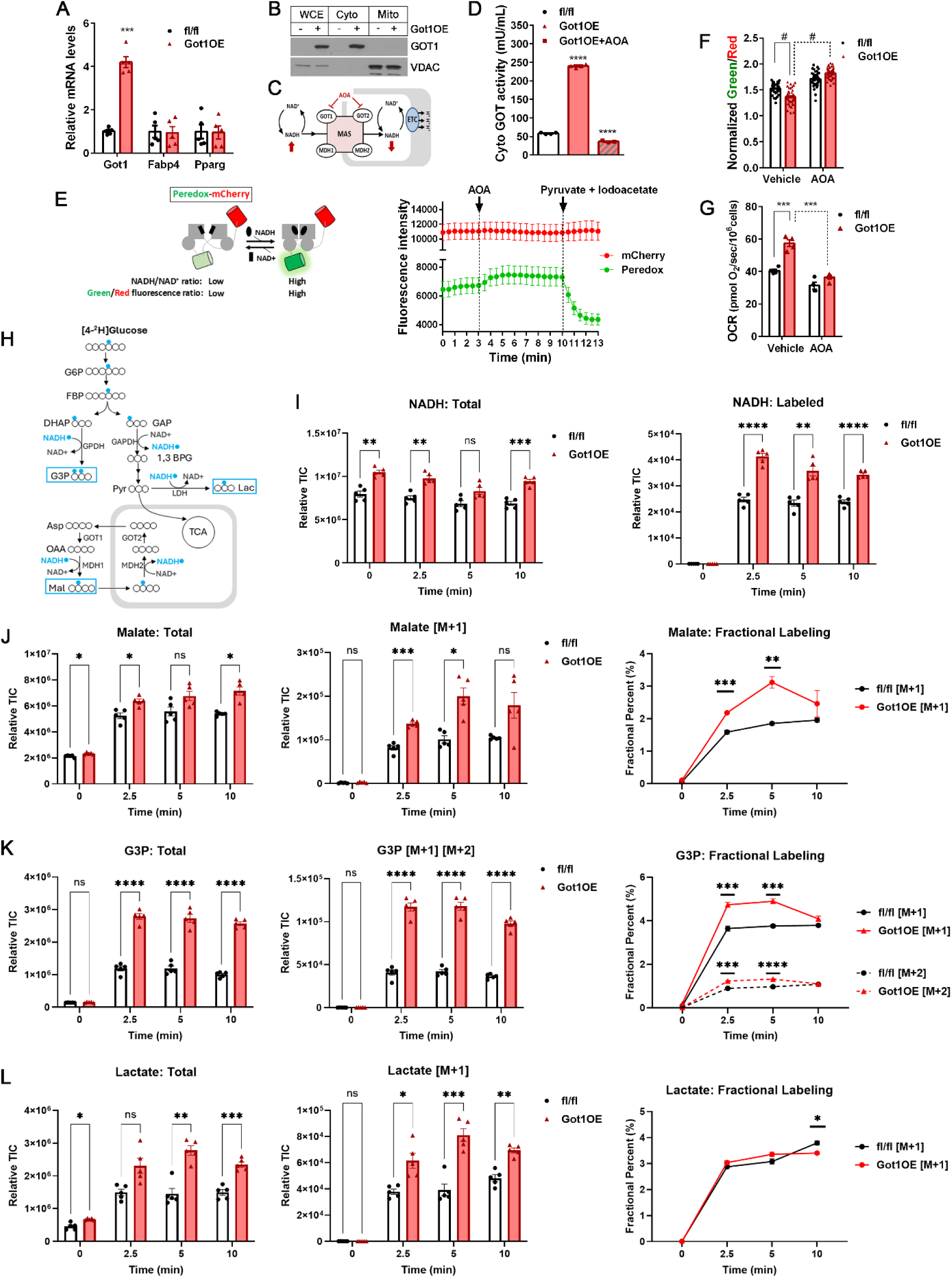
GOT1 overexpression activates the malate-aspartate shuttle in brown adipocytes. (A) qPCR analysis for adipogenesis in fully differentiated *Got1*fl/fl and *Got1*OE brown adipocytes. (B) WB analysis of GOT1 in the whole cell extracts, cytosolic, and mitochondrial fractions. (C) Schematic showing the effect of AOA-dependent inhibition of GOT1 and GOT2 in the MAS. Red arrows indicate the effect of AOA on NADH/NAD^+^ states. (D) Measurement of GOT1 enzyme activity in the cytosolic fraction of brown adipocytes treated with vehicle or 2 mM AOA. (E) Schematic showing Peredox-mCherry, a fluorescent sensor of the NADH-NAD^+^ redox state (left panel). A circularly permuted GFP T-Sapphire (green) is interposed between the two Rex subunits (gray), with an RFP mCherry (red) to normalize for the GFP fluorescence. NADH binding to Rex increases GFP fluorescence. A representative green and red fluorescence profile of Peredox-mCherry (right panel) prior to and after addition of 2 mM AOA, followed by addition of 20 mM pyruvate and 0.4 mM glycolytic inhibitor, iodoacetate. The Peredox signal was normalized to the lowest value with pyruvate and iodoacetate. (F) Effect of GOT1 overexpression on the cytosolic NADH-NAD^+^ redox state measured by the normalized green to red fluorescence ratio of Peredox-mCherry. Approximately 50 cells per group were analyzed. (G) Measurement of mitochondrial uncoupled respiration in brown adipocytes in the presence of an ATPase inhibitor, oligomycin. (H) Schematic of [^2^H] labeling used to determine cytosolic NADH-oxidizing pathways. The deuterium of NAD^2^H produced from [4-^2^H] glucose can be incorporated into malate, G3P, or lactate. (I) Total and [^2^H]-labeled NADH levels in *Got1*fl/fl and *Got1*OE brown adipocytes incubated with 25mM [4-^2^H] glucose at 2.5, 5, and 10 min. (J) Total and [M+1]-malate levels and fraction of [M+1]-malate after introducing [4-^2^H] glucose. (K) Total and [M+1]-, [M+2]-G3P levels and fraction of [M+1]- and [M+2]-G3P after introducing [4-^2^H] glucose. (L) Total and [M+1]-lactate levels and fraction of [M+1]-lactate after introducing [4-^2^H] glucose. All data are presented as the Mean ± SEM. \**p*<0.05, \*\**p*<0.01, \*\*\**p*<0.001, \*\*\*\**p*<0.0001, ^#^*p*<0.0001.

Thus, we monitored cytosolic NADH dynamics in *Got1*OE brown adipocytes by expressing a genetically encoded NADH biosensor Peredox ^50–53^. Peredox is constructed by linking a GFP variant to a bacterial NADH-binding protein, Rex (Fig. 4E, left panel). Upon binding NADH, Rex undergoes a conformational change that is coupled with an increase in green fluorescence. Peredox was tandemly fused to mCherry to enable ratiometric readout ^50^. The green and red fluorescence intensity was monitored over time in the same cell in the presence of glucose, followed by addition of AOA to inhibit GOT1 and of iodoacetate/pyruvate to inhibit glycolysis (Fig. 4E, right panel). GOT1 overexpression led to a lower green-to-red fluorescence ratio (Fig. 4F, vehicle), reflecting decreased NADH levels in the cytosol. This reduction was associated with an increase in mitochondrial respiration as measured by oxygen consumption rates (OCR) (Fig. 4G, vehicle), implying that reducing equivalents transferred from cytosolic NADH to the mitochondria via the MAS are fed into the ETC. Subsequent inhibition of GOT1 activity by AOA (Fig. 4C) rapidly increased the green-to-red fluorescence ratio in *Got1*OE brown adipocytes (Fig. 4F, AOA). In line with NADH accumulation in the cytosol, AOA abrogated the GOT1-dependent increase in mitochondrial respiration (Fig. 4G, AOA). These results clearly demonstrate that the overexpression of GOT1 in brown adipocytes activates MAS, which in turn facilitates the oxidation of cytosolic NADH to NAD^+^ with a concurrent transport of reducing equivalents to the mitochondria.

To further attest the GOT1-dependent activation of MAS in brown adipocytes, we performed stable isotope tracing using a [4-^2^H]-glucose tracer ^54–56^. Metabolism of [4-^2^H]-glucose in the glycolytic pathway produces [1-^2^H]-glyceraldehyde 3-phosphate (GAP) (Fig. 4H). GAPDH then transfers ^2^H from GAP to NADH. Cytosolic lactate dehydrogenase (LDH), malate dehydrogenase (MDH1), and glycerol-3-phosphate dehydrogenase (GPDH) subsequently transfer ^2^H from NAD^2^H to lactate, malate, and G3P, producing [2-^2^H]-lactate, [2-^2^H]-malate, and [2-^2^H]-G3P, respectively (Fig. 4H). GPDH can also produce [1, 2-^2^H]-G3P by transferring ^2^H from NAD^2^H to [1-^2^H]-dihydroxyacetone phosphate (DHAP). Thus, we assessed if GOT1 overexpression increases the production of deuterium-labeled malate by activating the MAS. Upon culturing brown adipocytes with [4-^2^H]-glucose, we detected rapid labeling of NADH from [4-^2^H]-glucose with higher levels of NAD^2^H in *Got1*OE brown adipocytes compared to control cells (Fig. 4I, right panel). The steady state isotopic labeling of metabolites was achieved as early as 2.5 minutes (Fig. 4J-4L). Indeed, GOT1 overexpression significantly increased the production of [M+1]-labeled malate (Fig. 4J, middle panel) and its labeled fraction (Fig. 4J, right panel), indicating that increased metabolic flux through GOT1 is sufficient to stimulate subsequent metabolic flux through the NADH-oxidizing enzyme MDH1 in the MAS pathway.

Accordingly, the combined results of Peredox, mitochondrial respiration, and [4-^2^H]-glucose tracing clearly demonstrate that GOT1 overexpression activates MAS, promoting the transport of reducing equivalents produced in glycolysis into mitochondria.

### GOT1-dependent activation of MAS enhances glucose uptake, glycolysis, and mitochondrial FA oxidation in brown adipocytes

Surprisingly, [M+1]- and [M+2]-labeled G3P (Fig. 4K, middle and right panels) and [M+1]-labeled lactate (Fig. 4L, middle panel) were also elevated in *Got1*OE brown adipocytes, suggesting that regeneration of cytosolic NAD^+^ via MAS may stimulate the glycolytic flux, which in turn leads to increased glucose uptake. To test this, we first assessed the glycolytic flux by measuring extracellular acidification ^22,57^ that is caused by the release of lactate produced during glycolysis. The extracellular acidification rates (ECAR) were indeed higher in *Got1*OE brown adipocytes compared to control cells (Fig. 5A). ECAR also remained higher during βAR stimulation with a βAR agonist, isoproterenol (ISO) (Fig. 5A). The inhibition of mitochondrial ATP synthase by oligomycin (Oligo) did not result in an additional increase in glycolysis, reflecting high rates of uncoupled respiration in brown adipocytes. Consistent with increased glycolysis, *Got1*OE brown adipocytes showed increased uptake of [^3^H]-2-DG, a non-metabolizable glucose analog (Fig. 5B, vehicle), coupled with elevated levels of GLUT1 glucose transporter (Fig. 5C). Glucose uptake also remained higher during βAR stimulation with isoproterenol (Fig. 5B). In line with a rise in glucose uptake and glycolysis, *Got1*OE brown adipocytes exhibited increased expression of genes involved in glucose transport and glycolysis (Fig. 5D). Intriguingly, despite the increase in glucose uptake and glycolysis, oxidation of the glycolytic end product pyruvate through the TCA cycle in the mitochondria was significantly diminished in *Got1*OE brown adipocytes treated with vehicle or isoproterenol (Fig. 5E). Instead, mitochondrial FA oxidation was greatly increased (Fig. 5F), in conjunction with elevated expression of FAO-related genes (Fig. 5D). *Got1*OE brown adipocytes exhibited higher mitochondrial respiration and oligomycin-independent leak respiration, which represents UCP1-mediated uncoupled respiration, and displayed a greater increase in total and leak respiration after treatment with isoproterenol (Fig. 5G). Enhanced uncoupled respiration was also accompanied by increased UCP1 levels in *Got1*OE brown adipocytes (Fig. 5I). Taken together, these data show that GOT1-dependent MAS activation stimulates glucose uptake and glycolysis while also increasing mitochondrial FA oxidation to support enhanced thermogenesis.

**Figure 5.**
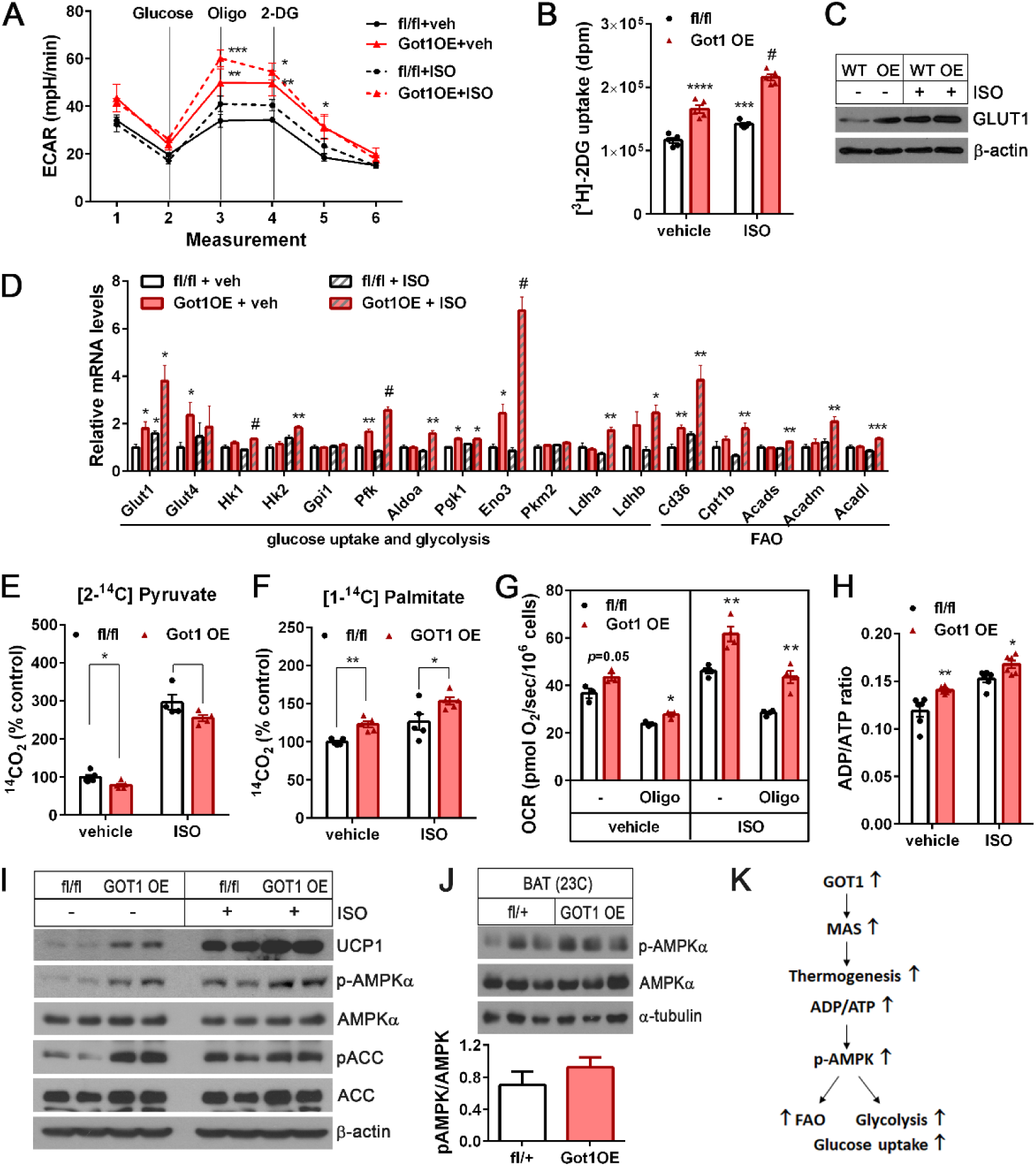
GOT1 overexpression enhances glucose uptake, glycolysis, and mitochondrial respiration in brown adipocytes. (A) Extracellular acidification rates (ECAR) measured by a Seahorse XF analyzer in *Got1*fl/fl and *Got1*OE brown adipocytes (n=10/group) at baseline and after addition of 10 mM glucose, followed by addition of 10 µM oligomycin and 50 mM 2-DG. (B) Measurement of [^3^H]-2DG uptake by *Got1*fl/fl and *Got1*OE brown adipocytes treated with vehicle or 5µM isoproterenol (ISO) for 4h. (C) WB analysis of GLUT1 protein expression. (D) qPCR analysis of genes involved in glucose uptake, glycolysis, and FA oxidation. (E, F) Oxidation of ^14^C-labeled pyruvate or palmitate in *Got1*fl/fl and *Got1*OE brown adipocytes treated with vehicle or isoproterenol for 4h. CO_2_ production was normalized by protein concentrations. (G) Mitochondrial respiration in brown adipocytes in the absence and presence of an ATPase inhibitor, oligomycin. (H) The ADP/ATP ratios measured in *Got1*fl/fl and *Got1*OE brown adipocyte lysates. (I, J) Increased phosphorylation of AMPKα at Thr172 in Got1-overexpressing brown adipocytes and BAT. (K) Schematic showing the effect of GOT1 overexpression on FAO, glucose uptake, and glycolysis. All data are presented as the Mean ± SEM. \**p*<0.05, \*\**p*<0.01, \*\*\**p*<0.001, ^#^*p*<0.0001.

Mitochondrial uncoupling has been shown to activate AMPK ^58–60^ due to the decrease in mitochondrial ATP production and a resulting increase in the ADP/ATP ratio, which subsequently stimulates adenylate kinase that converts two ADP into ATP and AMP ^61,62^. AMP binding to the γ subunit of AMPK promotes the phosphorylation of the α subunit at Thr172, enhancing AMPK’s catalytic activity ^63^. In fact, enhanced uncoupled respiration in *Got1*OE brown adipocytes resulted in a rise in the ADP/ATP ratio (Fig. 5H). Thus, we examined whether AMPK is activated in *Got1*OE brown adipocytes. The phosphorylation of AMPKα at Thr172 was increased in *Got1*OE brown adipocytes and the phosphorylated levels remained higher even after βAR stimulation, which activates AMPK ^64,65^ (Fig. 5I). Consistently, *Got1*OE BAT also exhibited a trend toward increased AMPKα phosphorylation at Thr172 (Fig. 5J). AMPK-mediated phosphorylation of acetyl-CoA carboxylase (ACC) at Ser79 has been shown to enhance FAO by increasing the activity of carnitine palmitoyl transferase (CPT1), the rate-limiting enzyme in FAO ^66^. In line with AMPK activation by phosphorylation at Thr172, ACC was highly phosphorylated at Ser79 in *Got1*OE brown adipocytes (Fig. 5I, vehicle), revealing that activated AMPK in part contributes to the observed increase in FAO. Given AMPK’s well-established role in promoting glucose uptake ^67–70^ and glycolysis ^71^, AMPK might also contribute to the observed increase in glucose uptake and glycolysis in *Got1*OE brown adipocytes.

Together, these data suggest that AMPK activation caused by enhanced uncoupled respiration is a feedback loop that signals a preference for FA in *Got1*OE brown adipocytes, while also supporting glucose uptake and glycolysis (Fig. 5K).

### Mice lacking Got1 in BAT have decreased capacity for FAO, but induce a compensatory increase in glucose oxidation to fuel cold-induced thermogenesis

To assess the importance of the cold-activated MAS in BAT thermogenesis, we generated BAT-specific *Got1* knockout (*Got1*^fl/fl:UCP1-cre/+^) mice by crossing *Got1*^fl/fl^ mice (EMMA, #10399) with *Ucp1*-Cre mice ^43^ (Fig. 6A). *Ucp1*-Cre mice express Cre recombinase mainly in BAT at room temperature and additionally in inguinal WAT (IWAT) undergoing brown remodeling during prolonged cold exposure ^43,72–75^. Cold-dependent induction of Got1 mRNA and protein in BAT and IWAT, but not in other tissues, was completely blunted in *Got1*^BATKO^ mice (Fig. 6B-C). However, the loss of GOT1 in BAT and IWAT depots had no impact on adipose remodeling during cold exposure (Fig. S1A), although it caused a reduction in the cellular NAD^+^/NADH ratio (Fig. 6H).

**Figure 6.**
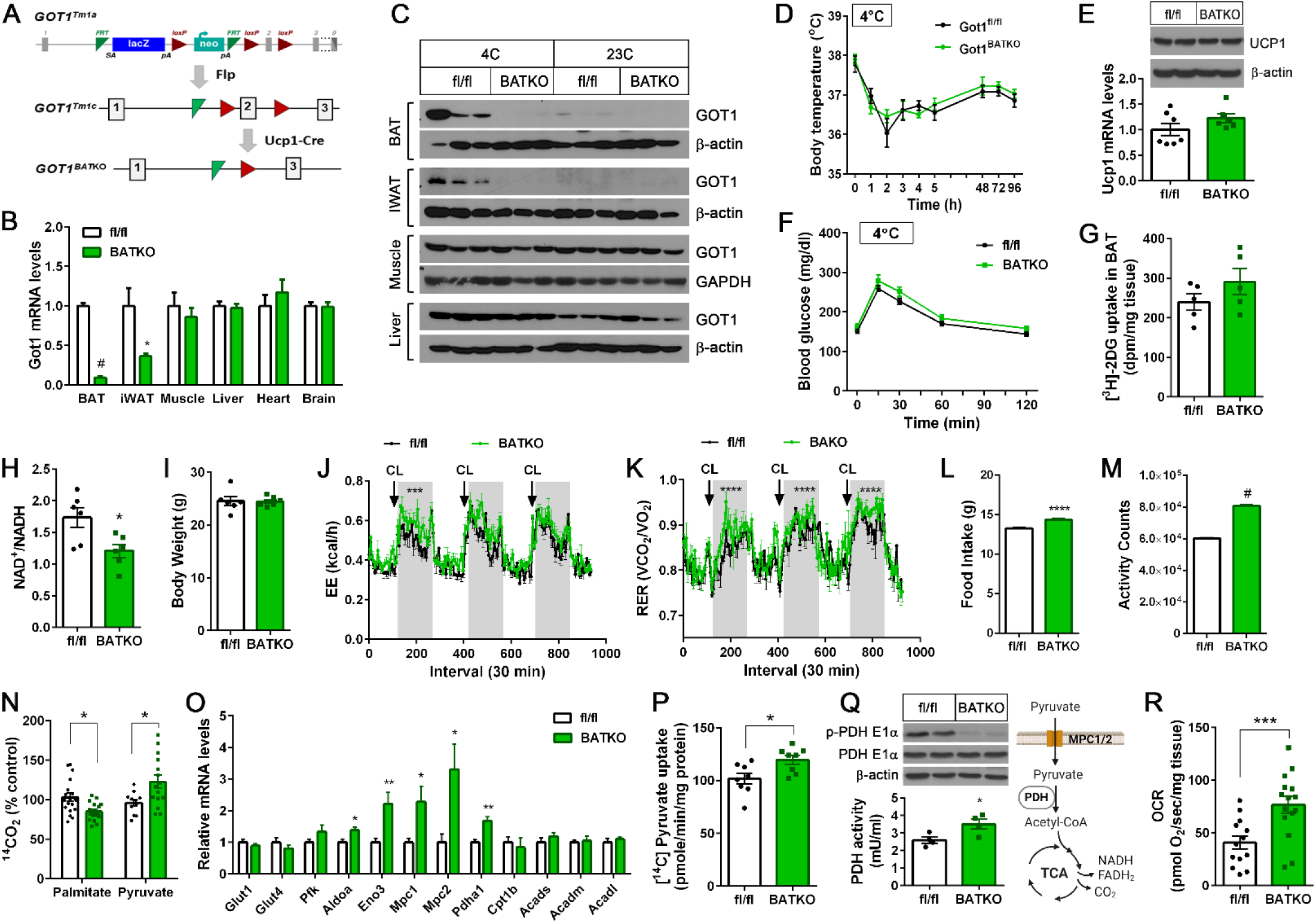
BAT-specific Got1 knockout mice exhibit reduced FA oxidation with a greater reliance on glucose oxidation for cold-induced thermogenesis. (A) Generation of BAT-specific *Got1* knockout mice. Exon 2 of the mouse *GOT1* gene was targeted for deletion by the LoxP/Cre system. (B) qPCR analysis for detection of *Got1* transcripts in *Got1*^fl/fl^ and *Got1*^BATKO^ male mice exposed to 4°C for 7 days (n=6-7/group). (C) WB analysis for GOT1 in *Got1*^fl/fl^ and *Got1*^BATKO^ mice housed at 23°C or exposed to 4°C for 7 days. (D) Body temperature of *Got1*^fl/fl^ and *Got1*^BATKO^ female mice during cold exposure (n=7-8/group). Core body temperature was measured using a rectal thermometer at indicated times. (E) WB and qPCR of UCP1 in BAT from the same mice described in D. (F) Glucose tolerance test of cold exposed-*Got1*^fl/fl^ and *Got1*^BATKO^ female mice (n=7-10/group). (G) Measurement of [^3^H]-2DG uptake by cold-activated BAT. (H) The NAD^+^/NADH ratio in BAT homogenates. (I-M) Measurement of body weight, energy expenditure, RER, food intake, and locomotor activity in *Got1*^fl/fl^ and *Got1*^BATKO^ male mice (n=6/group). Mice were placed in indirect calorimetric chambers and monitored during β-adrenergic stimulation of BAT with a β_3_AR agonist CL316243 (1mg/kg BW, daily). (N) Oxidation of ^14^C-labeled palmitate or pyruvate in BAT homogenates. BAT was extracted from mice exposed to 4°C for 7 days. (O) qPCR analysis of genes involved in glucose uptake, glycolysis, and FA oxidation in cold-activated BAT (n=6/group). (P) Uptake of ^14^C-labeled pyruvate by the mitochondria isolated from cold-activated BAT. (Q) Reduced phosphorylation of the PDH E1α subunit at Ser232 with a concurrent increase in PDH activity in Got1-deficient BAT. (R) Measurement of pyruvate-dependent mitochondrial respiration in BAT explants extracted from mice exposed to 4°C for 7 days. All data are presented as the Mean ± SEM. \**p*<0.05, \*\**p*<0.01, \*\*\**p*<0.001, \*\*\*\**p*<0.0001, ^#^*p*<0.0001.

We predicted that *Got1*^BATKO^ mice would be less cold-tolerant than control mice due to the impaired coordination of cold-stimulated FAO, glucose uptake, and glycolysis via MAS. However, contrary to our prediction, *Got1*^BATKO^ mice maintained their body temperature during cold exposure (Fig. 6D) along with normal levels of Ucp1 mRNA and protein in BAT (Fig. 6E). In addition, *Got1*^BATKO^ mice exhibited relatively normal glucose tolerance during GTT in the cold (Fig. 6F) with no significant difference in [^3^H]-2-DG uptake by BAT (Fig. 6G).

Indirect calorimetry further revealed that energy expenditure (EE) during BAT activation with a β_3_AR agonist CL316243 was comparable or slightly higher in *Got1*^BATKO^ mice compared to littermate controls (Fig. 6I-6J). Intriguingly, we noticed that *Got1*^BATKO^ mice had a greater reliance on glucose as a fuel source for BAT thermogenesis as evidenced by higher RER (Fig. 6K). The higher RER was coupled with an increase in food intake during indirect calorimetry (Fig. 6L). *Got1*^BATKO^ mice also showed increased locomotor activity (Fig. 6M), which may be related to the increased food intake event.

GOT1-deficient BAT extracted from cold-exposed *Got1*^BATKO^ mice had reduced capacity to oxidize FA in the mitochondria (Fig 6N). However, the oxidation of glucose (i.e. pyruvate) through the TCA cycle was significantly increased in *Got1*KO BAT (Fig. 6N). The increase in pyruvate oxidation coincided with upregulation of key genes encoding glycolytic enzymes (ALDOA, ENO), mitochondrial pyruvate carriers (MPC1/2), and pyruvate dehydrogenase (PDH) catalyzing the conversion of pyruvate to acetyl-CoA ^76,77^ (Fig. 6O). Moreover, the sequential pyruvate reactions, such as pyruvate import into the mitochondria (Fig. 6P), PDH enzyme activation by the reduced inhibitory phosphorylation (Fig. 6Q), and pyruvate-driven mitochondrial respiration (Fig. 6R), were all significantly accelerated in cold-activated *Got1*KO BAT. Likewise, *Got1*-deficient IWAT showed the increased expression of *Glut1*, *Glut4*, *Aldoa*, *Eno3*, and *Mpc1* genes compared to controls (Fig. S1B), suggesting that *Got1*-deficient beige adipocytes in cold-activated IWAT may undergo similar metabolic shifts in the absence of MAS activity. Together, these results establish that MAS activation by GOT1 is essential for promoting mitochondrial FA oxidation in BAT during cold exposure, and further suggest that impaired MAS activity rewires BAT mitochondria to import and oxidize more pyruvate to meet the energy demands for thermogenesis.

### Loss of MAS activity rewires mitochondrial fuel utilization toward glucose oxidation during βAR stimulation of brown adipocytes

To further attest the impacts of impaired MAS activity on mitochondrial fuel selection in brown adipocytes, we isolated stromal vascular fraction (SVF) from BAT of *Got1*^fl/fl^ mice, induced *Got1* deletion by retrovirus expressing Cre recombinase ^48^, and differentiated them into brown adipocytes followed by treatment with isoproterenol (ISO) for 4h. *Got1* deletion, confirmed by the loss of GOT1 expression and activity (Fig. 7A), did not alter brown adipogenesis as shown by no difference in adipogenic gene expression (Fig. 7B). First, we examined the effect of *Got1* deletion on MAS activity by utilizing an NADH biosensor Peredox and a [4-^2^H]-glucose tracer. As expected, *Got1*KO brown adipocytes expressing Peredox-mCherry exhibited an increased green to red fluorescence ratio compared to *Got1*fl/fl cells (Fig. 7C, vehicle), reflecting accumulation of NADH in the cytosol. Subsequent inhibition of ISO-induced GOT1 by AOA in *Got1*fl/fl brown adipocytes led to a rise in the green to red fluorescence ratio (Fig. 7C, AOA), but this effect was lost in *Got1*KO cells. The [4-^2^H]-glucose tracing (Fig. 7D) further confirmed the impaired MAS flux in *Got1*KO brown adipocytes stimulated with ISO. The production of deuterium-labeled malate (Fig. 7E) and its fractional enrichment (Supplementary Fig. 2A) were significantly diminished in *Got1*KO cells, indicating that hampered metabolic flux through GOT1 significantly impairs subsequent metabolic flux through MDH1 in the MAS pathway.

**Figure 7.**
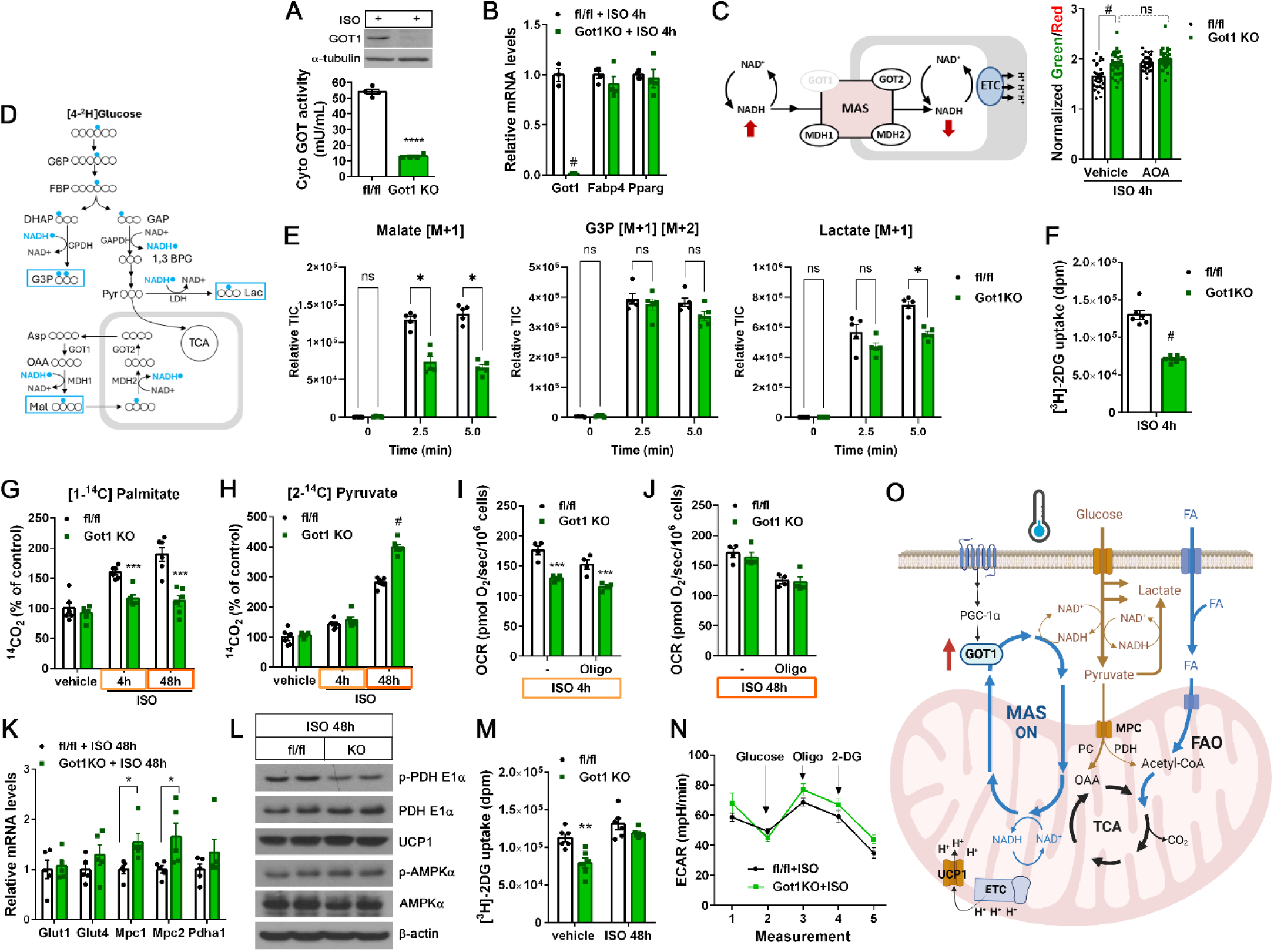
The loss of MAS activity by Got1 deletion diminishes mitochondrial FA oxidation during βAR stimulation, resulting in a compensatory increase in glucose oxidation. (A) Validation of loss of GOT1 expression and its enzymatic activity in the cytosolic fraction in *Got1*-deficient brown adipocytes treated with 5µM isoproterenol (ISO) for 4h. (B) qPCR analysis for adipogenic gene expression in *Got1*fl/fl and *Got1*KO brown adipocytes treated with ISO for 4h. (C) Schematic showing the effect of *Got1* deletion on the MAS-mediated NADH shuttling (left penal). Measurement of the cytosolic NADH-NAD^+^ redox state by Peredox-mCherry in *Got1*-deficient brown adipocytes (right panel). The green and red fluorescence signals of Peredox-mCherry were measured prior to and after addition of 2 mM AOA, followed by addition of 20 mM pyruvate and 0.4 mM glycolytic inhibitor, iodoacetate. (D) Schematic of [4-^2^H] glucose tracing. (E) [^2^H]-labeled malate, G3P, and lactate after introducing [4-^2^H] glucose to *Got1*fl/fl and *Got1*KO brown adipocytes treated with ISO for 4h. (F) [^3^H]-2DG uptake by brown adipocytes treated with ISO for 4h. (G, H) Oxidation of ^14^C-labeled palmitate or pyruvate in brown adipocytes treated with vehicle or ISO for 4h and 48h. (I, J) Mitochondrial respiration in brown adipocytes treated with ISO for 4h or 48h. (K) qPCR analysis of gene expression in brown adipocytes treated with ISO for 48h. (L) WB analysis of *Got1*fl/fl and *Got1*KO brown adipocytes treated with ISO for 48h. (M) [^3^H]-2DG uptake by brown adipocytes treated with vehicle or ISO for 48h. (N) Extracellular acidification rates (ECAR) in *Got1*fl/fl and *Got1*OE brown adipocytes (n=5/group) at baseline and after addition of 10 mM glucose, followed by addition of 10 µM oligomycin and 50 mM 2-DG. (O) A schematic diagram describing the role of GOT1 in cold-activated brown adipocytes. Cold-inducible GOT1 acts as a critical node that links cold-stimulated βAR signaling to the malate-aspartate shuttle (MAS). GOT1-dependent MAS activation facilitates the transport of reducing equivalents produced in glycolysis from the cytosol to mitochondria in the form of malate. This process is essential for the efficient oxidation of fatty acids (FA) to support thermogenesis under cold stress. Got1-deficient brown adipocytes rewire mitochondrial fuel preference toward glucose by increasing mitochondrial pyruvate import and PDH-mediated conversion of pyruvate to acetyl-CoA. All data are presented as the Mean ± SEM. \**p*<0.05, \*\**p*<0.01, \*\*\**p*<0.001, \*\*\*\**p*<0.0001, ^#^*p*<0.0001.

Interestingly, we noticed a reduced production of deuterium-labeled lactate, but not G3P, in *Got1*KO brown adipocytes (Fig. 7E). *Got1*KO brown adipocytes also took up less [^3^H]-2DG than *Got1*fl/fl cells (Fig. 7F), suggesting that impaired regeneration of cytosolic NAD^+^ from NADH by MAS reduces glycolytic flux and glucose uptake in *Got1*KO brown adipocytes. In addition, loss of MAS activity in *Got1*KO brown adipocytes resulted in an inability of mitochondria to enhance FA oxidation during βAR stimulation (Fig. 7G, ISO 4h). However, unlike the observed increase in glucose oxidation in *Got1*^BATKO^ mice, mitochondrial pyruvate oxidation was not upregulated in *Got1*KO brown adipocytes treated with ISO for 4h (Fig. 7H, ISO 4h). The resulting decrease in overall fuel supply to the ETC led to decreased mitochondrial respiration (Fig. 7I). These results clearly show that MAS plays a critical role in coordinating βAR-stimulated glucose uptake, glycolysis, and FAO in brown adipocytes. Next, we sought to test if increased pyruvate oxidation observed in *Got1*KO BAT occurs as a compensatory mechanism during prolonged βAR stimulation. To do this, we treated *Got1*fl/fl and *Got1*KO brown adipocytes with ISO for 48h. Indeed, pyruvate oxidation was significantly increased in *Got1*KO brown adipocytes (Fig. 7H, ISO 48h) while FAO remained decreased (Fig. 7G, ISO 48h). Moreover, the increased utilization of pyruvate as fuel for the ETC effectively restored mitochondrial respiration, enhancing both total and uncoupled respiration in *Got1*KO brown adipocytes treated with ISO for 48h (Fig. 7J). These results thus indicate that *Got1*KO brown adipocytes compensate for reduced FAO by rewiring mitochondrial fuel utilization toward pyruvate oxidation during prolonged βAR stimulation. This metabolic shift was also associated with upregulation of *Mpc1* and *Mpc2* gene expression (Fig. 7K) and reduction of inhibitory phosphorylation of PDH (Fig. 7L), as seen in BAT of cold-exposed *Got1*^BATKO^ mice. In line with increased glucose utilization for mitochondrial respiration, glucose uptake was restored to normal levels in *Got1*KO brown adipocytes treated with ISO for 48h (Fig. 7M), with glycolysis being relatively comparable (Fig. 7N). Under βAR stimulation that activates AMPK ^64,65^, AMPKα phosphorylation at Thr172 was not different between *Got1*fl/fl and *Got1*KO brown adipocytes (Fig. 7L), indicating that AMPK has no effect on these changes. ECAR (Fig. 7N) suggests that LDH helps NAD^+^ regeneration for glycolysis in the absence of MAS. Taken together, the compensatory mechanisms occurring both in the cytosol and mitochondria during prolonged βAR stimulation likely explain the cold tolerance and glucose tolerance phenotypes of *Got1*^BATKO^ mice exposed to cold (Fig. 6D and 6F).

## Discussion

During cold adaptation, BAT simultaneously metabolizes both fatty acids (FA) and glucose but distinctly utilizes them; FA is primarily used as a fuel source for thermogenesis, whereas glucose-derived carbons are mostly released as lactate or channeled into various biosynthetic pathways. In the present study, we show that the malate-aspartate shuttle (MAS) is essential for sustaining mitochondrial FA utilization in BAT under cold conditions. Intriguingly, MAS activity in BAT is tightly regulated by environmental temperatures through drastic changes in GOT1 protein levels. At room temperature, GOT1 is present at barely detectable levels in BAT, but its expression is significantly induced by cold via the βAR-cAMP-PKA-PGC-1α/NT-PGC-1α axis. This regulatory mechanism of GOT1 is unique in BAT, as GOT1 is abundantly expressed in many other tissues. Using BAT-specific *Got1* knock-in and *Got1* knockout mice and their respective brown adipocytes, we have clearly demonstrated that cold-inducible GOT1 is a molecular switch turning on the MAS, enabling BAT to increase FA oxidation rates to fuel thermogenesis under cold conditions (Fig. 7O).

How does MAS promote mitochondrial FA oxidation in BAT? MAS oxidizes glycolysis-derived NADH to NAD^+^ in the cytosol, maintaining the high cytosolic NAD^+^/NADH ratio required for continued glycolysis. We found that GOT1 overexpression in brown adipocytes increases the expression of genes linked to FA uptake (*Cd36*), mitochondrial transport (*Cpt1b*), and FA β-oxidation (*Acads*, *Acadm*, A*cadl*). When the cytosolic NAD^+^/NADH ratio is high, NAD^+^-dependent SIRT1 deacetylase has been shown to promote transcription of FAO-related genes, including *Cpt1b*, *Acadm*, and *Acadl*, by facilitating the interaction of PGC-1α with PPARα ^78–80^. Thus, our findings suggest that the high cytosolic NAD^+^/NADH ratio maintained by MAS could contribute to enhanced expression of FAO-related genes in BAT under cold conditions. The oxidation of acetyl-CoA derived from FA within the TCA cycle is initiated by combining with oxaloacetate (OAA). A recent glucose tracing study ^23^ showed that cold-activated BAT increases the flux of pyruvate to OAA via pyruvate carboxylase (PC) over acetyl-CoA via PDH, suggesting that glucose supplies OAA necessary for the efficient oxidation of FA-derived acetyl-CoA through the TCA cycle. This is consistent with previous findings that acetyl-CoA activates PC but inhibits PDH ^81^. MAS transports electrons derived from cytosolic NADH into mitochondria in the form of malate, which is oxidized back to OAA, regenerating NADH in the mitochondria. It should also be noted that MAS is interconnected with the TCA cycle through MDH2 and shared intermediates (e.g., malate and OAA). By shuttling reducing equivalents and supporting anaplerotic reactions, MAS may enable BAT to efficiently handle the high flux of acetyl-CoA from FA β-oxidation without depleting TCA intermediates.

Alternatively, given MAS’s role in exporting mitochondrial OAA to the cytosol in the form of aspartate, MAS may support the enhanced flux of PC over PDH in cold-activated BAT by clearing excess OAA derived from pyruvate out of the mitochondria. Although the precise mechanism remains to be clarified, our data show that impaired MAS activity by GOT1 deficiency reduces the ability of BAT mitochondria to efficiently oxidize FA under cold stress, leading to a compensatory increase in mitochondrial pyruvate import, its conversion to acetyl-CoA via PDH, and subsequent oxidation through the TCA cycle.

GOT1 overexpression in brown adipocytes enhanced glucose uptake and glycolysis in the cytosol, whereas GOT1 deficiency led to a significant decrease in these processes under β-AR stimulation. Interestingly, reduced glucose uptake and glycolysis in *Got1*-deficient brown adipocytes were restored during prolonged β-AR stimulation, coinciding with increased utilization of glucose as fuel for thermogenesis. These findings indicate that other cytosolic NADH-oxidizing pathways, such as lactate dehydrogenase (LDH), a glycerol-3-phosphate (G3P) shuttle composed of cytosolic and mitochondrial G3P dehydrogenases ^82^, and a recently identified AIFM2 NADH oxidase ^83^, could compensate for the loss of MAS in the cytosol to support glycolysis. However, despite their ability to replenish cytosolic NAD^+^ from NADH, these NADH-oxidizing pathways use different mechanisms to transfer electrons derived from cytosolic NADH, reflecting each pathway serving different purposes based on tissue-specific metabolic demands and cellular requirements. LDH transfers electrons to lactate, which is released to the bloodstream. The G3P shuttle and AIFM2 transfer electrons directly to the ubiquinone (Q) molecule within the ETC ^82,83^. However, MAS transports electrons across the IMM by leveraging a series of coordinated reactions and metabolite exchanges between the cytosol and mitochondria. Our findings suggest that this distinct mechanism of MAS is essential for sustaining FA oxidation in BAT mitochondria during increased thermogenic demand.

In summary, the present study identifies brown adipocyte GOT1 as a critical node that links cold-stimulated β-adrenergic signaling to the MAS, leading to efficient FA oxidation to produce energy for thermogenesis. This discovery expands our understanding of MAS beyond redox balance and electron transfer, positioning it as a critical regulator of mitochondrial fuel selection in BAT.

### Limitations of the study

Although the present study has shown that GOT1-dependent MAS activation is crucial for sustaining mitochondrial FA oxidation in BAT, we cannot exclude the possibility that the non-MAS function of GOT1 indirectly contributes to enhanced FA oxidation in BAT. Given GOT1’s role in converting aspartate to OAA, GOT1 may supply OAA to PEPCK-C in glyceroneogenesis ^84^, a metabolic pathway creating G3P from non-glucose precursors for replenishment of intracellular TAG, although quantitative significance of glyceroneogenesis in cold-activated BAT is not clear. A recent report ^85^ suggests that the widely used *Ucp1*-Cre line ^43^ has leaky Cre expression in various non-adipose tissues. To mitigate the potential confounding effects on metabolic readouts, we utilized a β_3_-AR agonist CL316243 to selectively activate BAT in R26-*Got1*^BATOE/+^ and *Got1*^BATKO^ mice, in conjunction with parallel cell culture studies using their respective brown adipocytes. Finally, it remains to be elucidated whether MAS plays a similar role in human brown fat, as the *Got1* gene is highly expressed in cold-activated human brown fat as well ^34^.

## Resource availability

### Lead contact

Further information and requests for resources and reagents should be directed to and will be fulfilled by the Lead Contact, Ji Suk Chang.

### Materials availability

All unique reagents generated in this study are available from the lead contact with a completed Materials Transfer Agreement.

## Acknowledgements

We thank Dr. Tom Gettys (Pennington Biomedical Research Center) for providing PGC-1α and UCP1 antibodies, Ms. Jisu Lee for technical assistance, and Drs. Randall L Mynatt and Jingying Zhang (PBRC Transgenics Core) for technical support in generating the Got1-knockin mouse line. This work was supported by the National Institutes of Health grant R01DK136536 (JSC) and used Cell Biology & Bioimaging Core, Genomics Core, and Animal Metabolism & Behavior Core that are supported in part by COBRE (1P20GM135002-01) and NORC (P30DK072476) center grants from the National Institutes of Health.

## Author Contributions

C.H.P. conducted experiments, analyzed data, and interpret data. M.P., M.E.K., and H.C. conducted experiments and analyzed data. S.R.L. generated Got1 knock-in mice. C.J. analyzed data. J.S.C. conceived of the presented idea, interpreted data, and wrote the manuscript. J.S.C. is the guarantor of this work and, as such, had full access to all the data in the study and takes responsibility for the integrity of the data and the accuracy of the data analysis. All authors read and approved the submitted version.

## Supplemental Information

Figures S1-S2

## Methods

### Experimental model and study participant details Mice

Gene targeting and Got1 knock-in mouse production were performed by the Transgenics Core at Pennington Biomedical Research Center. DNA derived from a C57BL6 genomic BAC library clone (RP24-399J12) was used to generate a targeting vector containing the CAG-loxP-neo-3xpA(stop)-loxP-Got1. Targeted albino B6 embryonic stem cells carrying the transgene at the Rosa26 locus were injected into C57BL6 blastocytes, and chimeric mice were mated to C57BL6 mice to obtain heterozygous offspring containing a germline transgene allele. Heterozygous R26-*Got1*^fl/+^ mice were then mated to obtain homozygous R26-*Got1*^fl/fl^ mice. To generate brown adipose tissue (BAT)-specific Got1 knock-in mice, R26-*Got1*^fl/fl^ mice were mated to *Ucp1*-Cre transgenic mice ^43^ (Jackson Laboratory, #024670) that express Cre recombinase under the control of *UCP1* promoter, which is activated in brown adipocytes. While the polyadenylation sequence terminates transcription initiated by the CAG promoter, LoxP/Cre-mediated excision of the stop cassette enables targeted expression of *Got1* in BAT in R26-*Got1*^fl/+:Ucp1-Cre/+^ mice. For generation of BAT-specific *Got1* knockout mice, C57BL/6N-*Got1*^tm1a(EUCOMM)Hmgu^/H mice (EMMA, #10399) containing an FRT-flanked lacZ/neomycin cassette followed by a loxP site upstream of *Got1* exon 2 were first mated to FLP mice to remove the lacZ and neo cassettes. The resulting Got1^fl/fl^ mice were mated to *Ucp1*-Cre transgenic mice to generate *Got1*^fl/+:Ucp1-Cre/+^ mice, which were then mated with *Got1*^fl/fl^ mice to generate homozygous *Got1*^fl/fl:Ucp1-Cre/+^ mice. Mice were housed at room temperature of 23°C under a 12-h light/12-h dark cycle with ad libitum feeding (standard chow 5001, LabDiet, St. Louis, MO). Animal studies were carried out in accordance with the institutional guidelines and were approved by the Institutional Animal Care and Use Committee of the Pennington Biomedical Research Center. 10-to-19-week-old male and female mice were used for animal experiments.

### Brown adipocytes

Stromal vascular fraction (SVF) containing brown preadipocytes was isolated from interscapular BAT of newborn R26-*Got1*^fl/fl^ or *Got1*^fl/fl^ pups by collagenase digestion and immortalized by infection with SV40T antigen-expressing retrovirus as previously described ^35^. LoxP/Cre-mediated overexpression or deletion of *Got1* was then induced by retrovirus expressing Cre recombinase (AddGene) ^48^. The resulting *Got1*-overexpressing or -deficient brown preadipocytes were grown to confluence in culture medium supplemented with 20 nM insulin and 1 nM T3 (differentiation medium) and induced for differentiation by incubating in differentiation medium supplemented with 0.5 mM isobutylmethylxanthine (IBMX), 0.5 µM dexamethasone, and 0.125 mM indomethacin for 48 hours, as previously described ^86^. Thereafter, the cells were maintained in differentiation medium until day 7. Fully differentiated brown adipocytes were treated with or without 5 µM of a β-adrenergic receptor agonist isoproterenol to stimulate β-AR signaling.

### Method details Cold tolerance tests

Mice were singly housed and exposed to 5°C for 6h or for 7 to 10 days. Core body temperature was measured at baseline and during cold exposure using a rectal probe RET-3 (a thin stainless-steel shaft 19 mm long with a smooth ball tip of 1.7 mm diameter).

### Indirect calorimetry

For metabolic phenotyping, mice were placed in indirect calorimetry chambers (Sable Systems International, North Las Vegas, NV) and monitored for energy expenditure (kcal/h), VO_2_ and VCO_2_ at 28°C prior to and after intraperitoneal injections of a β_3_-adrenergic receptor agonist, CL316243 (1 mg/kg body weight, daily). Food intake and locomotor activity were also measured while the mice were in the chambers.

### Glucose tolerance tests

Mice housed at room temperature were fasted for 16 h and injected intraperitoneally with a glucose bolus (2 g/kg body weight). Mice exposed to 5°C were fasted for 5 h and injected with a glucose bolus (2 g/kg body weight). Blood glucose levels were measured using a Contour Next EZ glucometer (Bayer, Leverkusen, Germany).

### ChIP assay

Chromatin immunoprecipitation was performed as described previously ^33,36^. Briefly, BAT explants were chopped into small pieces and fixed with paraformaldehyde. After incubation for 10 min, 2.5M glycine was added to a final concentration of 0.125 M and the material was pelleted by centrifugation at 8,000 rpm and resuspended in ChIP lysis buffer (10 mM Tris-HCl, pH 8.0, 10 mM NaCl, 3 mM MgCl_2_, 0.5% (vol/vol) NP-40, protease inhibitors). The homogenates were centrifuged at 1,200 × g and the nuclear pellets were sonicated using a Bioruptor in ChIP shearing buffer (0.25% SDS, 10 mM Tris-HCl, pH 8.0, 1 mM EDTA, protease inhibitors). Immunoprecipitation was conducted with rabbit polyclonal PGC-1α antibody ^35,41^ directed against the N-terminus of PGC-1α. DNA-protein complexes were eluted from protein A beads with elution buffer (100 mM NaHCO_3_, 1% SDS) and reverse crosslinked by adding NaCl to a final concentration of 0.2 M and incubating at 65°C. PCR was performed to determine the recruitment of PGC-1α/NT-PGC-1α to the *GOT1* promoter region using the following primers: for ERRE1 site, forward 5′-CTAATCCCCACAACAACCTG, reverse 5′-CTGTCCAGATCCTTTCTCCA, for ERRE2 site, forward 5′-CAAGGTCCTTTAGGGCTAGG, reverse 5′-TGGCACAGGACTCTAGTGGT.

### Measurement of cytosolic NAD^+^/NADH ratios

Cytosolic NAD^+^/NADH ratio was measured using a genetically encoded fluorescent NADH biosensor Peredox-mCherry (Addgene, #32383) ^50^. Brown adipocytes were incubated in phenol red-free DMEM containing 0.5 mM glucose and 1% fatty acid-free BSA and imaged by a Lionheart FX Automated Microscope (BioTek) with a 10× objective in an environmentally controlled chamber at 37 °C and 5% CO2. Green and red fluorescence images were acquired at 400 nm excitation/525 nm emission and 586 nm excitation/647 nm emission every 30 s for 15 min, during which time 2 mM aminooxyacetate (AOA) and then 0.4 mM Iodoacetate/20 mM pyruvate were added to inhibit GOT activity and glycolysis, respectively. Using ImageJ software, images were background corrected, and a pixel-by-pixel green-to-red ratio image was generated for each time point as described previously ^50^. Total intracellular NAD^+^/NADH ratio was measured in BAT tissue homogenates using an EnzyChrom NAD^+^/NADH assay kit (BioAssay Systems).

### Mitochondrial respiration assay

Oxygen consumption rates (OCR) of brown adipocytes and BAT explants were measured as described previously ^41^. Briefly, cultured brown adipocytes (10^6^ cells) or fresh BAT explants (5 mg) were placed in a magnetically stirred respirometric chamber of the OROBOROS Oxygraph-2k (Oroboros Instruments, Innsbruck, Austria) containing the culture medium or respiration buffer ^41^. For cultured brown adipocytes, OCR measurements were obtained at baseline and after injection of oligomycin and antimycin A. For BAT explants, OCR was measured in the presence of malate/pyruvate, followed by injection of antimycin A. The value of basal and leak mitochondrial respiration was determined by subtracting antimycin A-independent non-mitochondrial respiration as described in the Oroboros Operator’s Manual.

### Stable isotope labeling and LC-MS analysis

A [4-^2^H]-glucose tracer was utilized to label glycolysis-derived NADH with deuterium and trace the flow of NAD^2^H to metabolites generated by NADH-dependent dehydrogenase activity as previously described ^54–56^. For isotopic labeling, brown adipocytes were cultured in 6-well plate in phenol red-, glucose-, pyruvate-, and glutamine-free DMEM supplemented with 25 mM [4-^2^H]-glucose, 2 mM L-glutamine, and 10% dialyzed FBS. At 2.5-, 5-, and 10-min time points, cells were washed three times in 0.9% saline and immediately quenched in 1 ml of pre-cooled 80% methanol containing 1 µM of ^15^N-valine as an internal control. After freezing at −80°C for 15 min, cells were scrapped and cleared by centrifugation at 4°C. To measure soluble metabolites in cell extracts, 15 μl supernatant was injected into LC-MS. A quadrupole orbitrap mass spectrometer (Q Exactive; ThermoFisher Scientific) operating in negative or positive ion mode was coupled to a Vanquish UHPLC system (ThermoFisher Scientific) with electrospray ionization and used to scan from m/z 80 to 680 at 2 Hz, with a 140,000 resolution. LC separation was achieved on an XBridge BEH Amide column (2.1 x 150 mm^2^, 2.5μm particle size, 130 Å pore size; Waters Corporation) using a gradient of solvent A (95:5 water: acetonitrile with 20 mM of ammonium acetate and 20 mM of ammonium hydroxide, pH 9.45) and solvent B (acetonitrile). Flow rate was 150μl/min. The LC gradient was: 0 min, 85% B; 2 min, 85% B; 3 min, 80% B; 5 min, 80% B; 6 min, 75% B; 7 min, 75% B; 8 min, 70% B; 9 min, 70% B; 10 min, 50% B; 12 min, 50% B; 13 min, 25% B; 16 min, 25% B; 18 min, 0% B; 23 min, 0% B; 24 min, 85% B; and 30 min, 85% B. The autosampler temperature was 5°C. LC-MS data was converted from .raw files to .mzXML and then loaded into EI-MAVEN open source software ^87^ (Version 0.12.0, Elucidata). Using our annotated MS library, containing retention times and accurate m/z values, we selected metabolite peaks of interest on EI-MAVEN. Peak shape and quality were evaluated and Area Top of the metabolites of interest were recorded. Natural isotope correction was performed with AccuCor2 R code (https://github.com/wangyujue23/AccuCor2) and IsoCorrectoR ^88^. Microsoft Excel and R were used for data analysis.

### 14C-palmitate and ^14^C-pyruvate oxidation assay

Palmitate oxidation was measured as previously described ^41,89^. In brief, freshly isolated BAT explants were homogenized in SET buffer containing 250 mM sucrose, 1 mM EDTA, and 10 mM Tris-HCl (pH 7.4). Tissue homogenates were immediately incubated for 30 min at 37 °C with the reaction mixture ^89^ containing 200 µM palmitic acid coupled to BSA and [1-^14^C]-palmitate (0.5 µCi/ml). At the end of incubation, 70% perchloric acid was added to stop the reaction. ^14^CO_2_ produced during the incubation was trapped in 1M NaOH and the acidified portion of the incubation mixture was collected for liquid scintillation counting in a Perkin Elmer Tri-Carb 2910TR liquid scintillation analyzer. Pyruvate oxidation was determined as above by incubating tissue homogenates in the reaction mixture containing [2-^14^C]-pyruvate (0.5 µCi/ml), 1 mM pyruvate, and 1 mM thiamine pyrophosphate. Total palmitate/pyruvate oxidation was normalized to protein content in each sample.

### Mitochondrial pyruvate uptake assay

Mitochondrial pyruvate uptake assay was performed as previously described ^90^. Briefly, freshly isolated BAT mitochondria were resuspended to 15.0 mg/ml in Uptake buffer (120 mM KCl, 5 mM HEPES, pH 7.4, 1 mM EGTA, 2 µM rotenone, and 2 µM antimycin A). The mitochondria were split into two equal volumes and treated for 5 min on ice with or without 2 mM α-Cyano-4-hydroxycinnamic acid (CHC) to inhibit mitochondrial pyruvate carrier (MPC). To start the pyruvate uptake assay, 35 μl of CHC-treated or untreated mitochondria were mixed with an equal volume of 2× Pyruvate buffer (120 mM KCl, 5 mM HEPES, pH 6.1, 1 mM EGTA, 2 µM rotenone, and 2 µM antimycin A, and 100 µM [2-^14^C]-pyruvate). After 30 seconds, the uptake was stopped by adding 175 μl of Stop buffer (108 mM KCl, 4.5 mM HEPES, pH 6.8, 0.9 mM EGTA, 1.8 µM rotenone, 1.8 µM antimycin A, and 4 mM CHC). After stopping the reaction, the mitochondria were separated from the reaction mixture by centrifugation at 10,000× g, washed twice, and collected for liquid scintillation counting. Counts from CHC-treated mitochondria (negative control) were subtracted from untreated mitochondria.

### Glucose uptake assay

Brown adipocytes were pre-incubated in serum-free DMEM supplemented with 1% fatty acid-free BSA. After rinsing with PBS, the cells were incubated for 10 min in glucose-free DMEM supplemented with 1 µCi of D-[^3^H]-2-deoxyglucose (2-DG), 0.2 mM of 2DG, and 1% fatty acid-free BSA, in the presence and absence of 20 µM cytochalasin B. Following incubation, 0.2 mM cytochalasin B was added to terminate glucose transport. The cells were then washed three times with ice-cold PBS and collected for liquid scintillation counting. Counts from cytochalasin B-pretreated cells (negative control) were subtracted from untreated cells. Ex vivo glucose uptake assay was performed as above with a slight modification. For glucose transport, freshly isolated BAT explants (approximately 30 mg) were incubated for 20 min in Krebs-Ringer bicarbonate buffer (KRBB) supplemented with 1 µCi of [^3^H]-2-DG and 0.2 mM of 2DG, in the presence and absence of cytochalasin B. Total glucose uptake was normalized to tissue weight.

### Measurement of extracellular acidification rate (ECAR)

To monitor rates of glycolysis, extracellular acidification caused by lactate produced during glycolysis was measured as previously described ^91^. Briefly, brown adipocytes were plated in Seahorse XF 24-well plates and equilibrated in serum-free Seahorse XF media at 37°C for 45 min. ECAR was measured at baseline and after sequential addition of 10 mM glucose, 10 µM oligomycin, and 50 mM 2-DG in accordance with manufacturer’s instructions.

### Enzyme activity assay

Enzyme activity of glutamic-oxaloacetic transaminase (GOT) and pyruvate dehydrogenase (PDH) was measured in BAT tissue homogenates by the GOT and PDH Activity Assay kits (Sigma).

### Measurement of the ADP/ATP ratios

The cellular ADP/ATP ratios were measured by the ADP/ATP Ratio Assay kit (Sigma).

### Western blot analysis

Whole cell extracts were prepared from tissues or brown adipocytes by homogenization in lysis buffer ^86^ in the presence of a protease and phosphatase inhibitor cocktail (Roche). Lysates were separated by SDS-PAGE, transferred to nitrocellulose membrane, and analyzed by immunoblotting with following antibodies: GOT1 antibody (Pro Sci, #30-379), GOT2 antibody (Invitrogen, #PA5-27572), MDH1 antibody (Santa Cruz, #sc-166879), MDH2 antibody (Invitrogen, #PA5-21700), UCP1 antibody ^92^, GAPDH antibody (Abcam, #ab9485), VDAC antibody (Abcam, #ab15895), α-tubulin antibody (Abcam, #ab7291), β-actin antibody (Sigma, #A5441), phospho-PDH E1α antibody (S232) (Millipore, #AP106350). PDH E1α antibody (#3205), MPC1 antibody (#14462), MPC2 antibody (#46141), GLUT1 antibody (#12939), phospho-AMPKα (T172) antibody (#2531L), AMPKα antibody (#2532), phospho-ACC (S79) antibody (#3661), and ACC antibody (#3662) were purchased from Cell Signaling.

### Quantitative real-time PCR analysis

Total RNA was reverse transcribed for quantitative real-time PCR analysis as described previously ^36,86^. Gene expression analysis was performed using the Applied Biosystems 7900 (Applied Biosystems) and iTaq Universal SYBR Green Supermix (Bio-Rad). Relative mRNA expression of the genes of interest was determined using gene-specific primers after normalization to cyclophilin by the 2^-ΔΔCt^ method. The validated primer sequences were obtained from PrimerBank public resource ^93^.

### Histological analysis

Tissue samples were fixed in 10% neutral-buffered formalin, paraffin embedded, and sectioned (5 µm) by the Cell Biology & Bioimaging Core at Pennington Biomedical Research Center.

Hematoxylin and eosin-stained paraffin sections were scanned using a Hamamatsu NanoZoomer slide scanner (Hamamatsu, Japan).

### Statistical analysis

All line and bar graphs were created by using Prism 10 software (GraphPad Software, San Diego, CA, USA) and student *t* test or two-way ANOVA was used to compare the differences between groups. Data are presented as mean ± SEM. Values of *P* < 0.05 were considered statistically significant.

---

**Figure S1.**
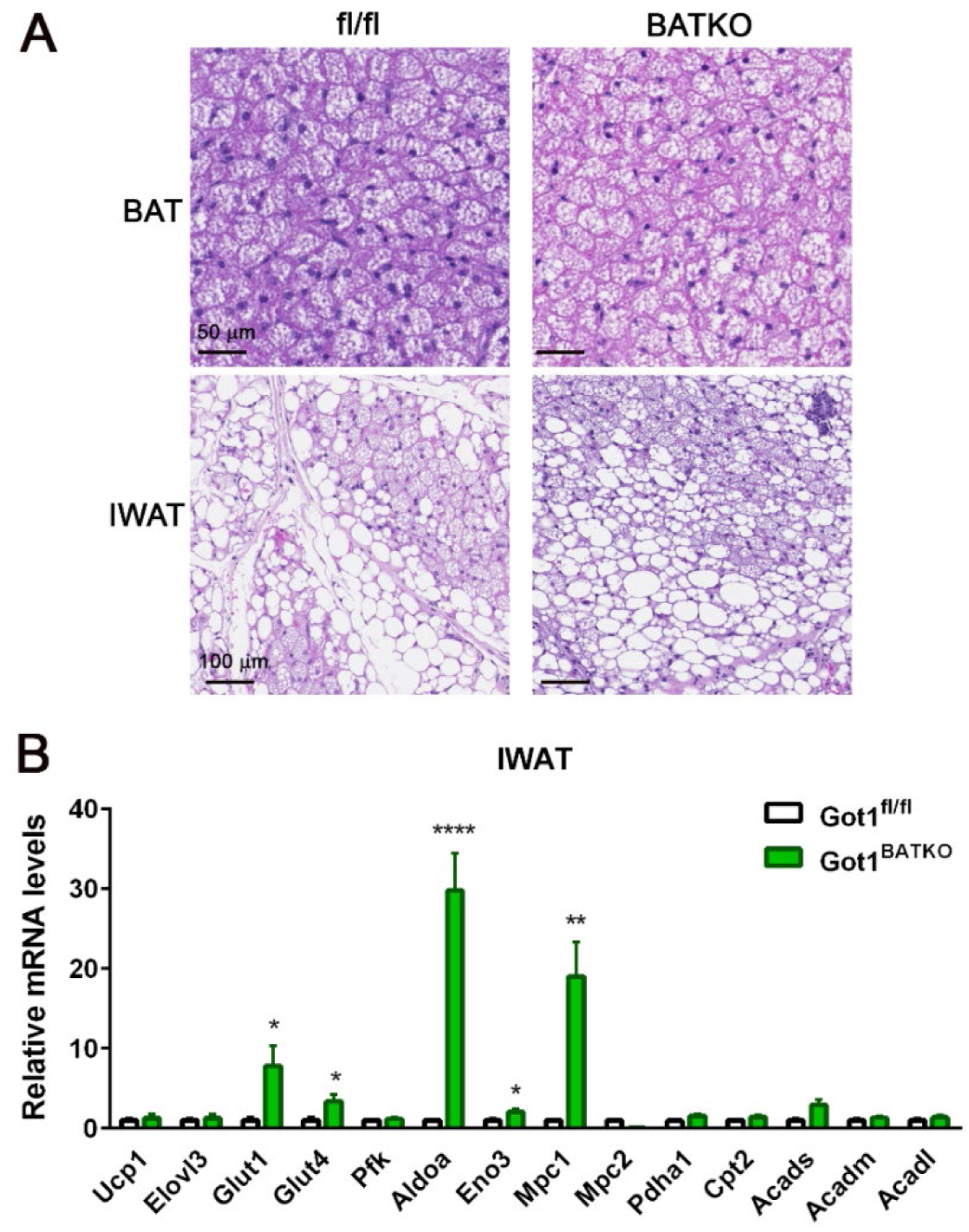
BAT-specific Got1 knockout mice exhibit normal adipose remodeling during prolonged cold exposure. (A) H&E staining analysis of BAT and IWAT from *Got1*^fl/fl^ and *Got1*^BATKO^ female mice exposed to 4°C for 10 days. (B) qPCR analysis of genes involved in glucose uptake, glycolysis, and FA oxidation in cold-activated IWAT (n=6/group). All data are presented as the Mean ± SEM. \**p*<0.05, \*\**p*<0.01, \*\*\*\**p*<0.0001.

**Figure S2.**
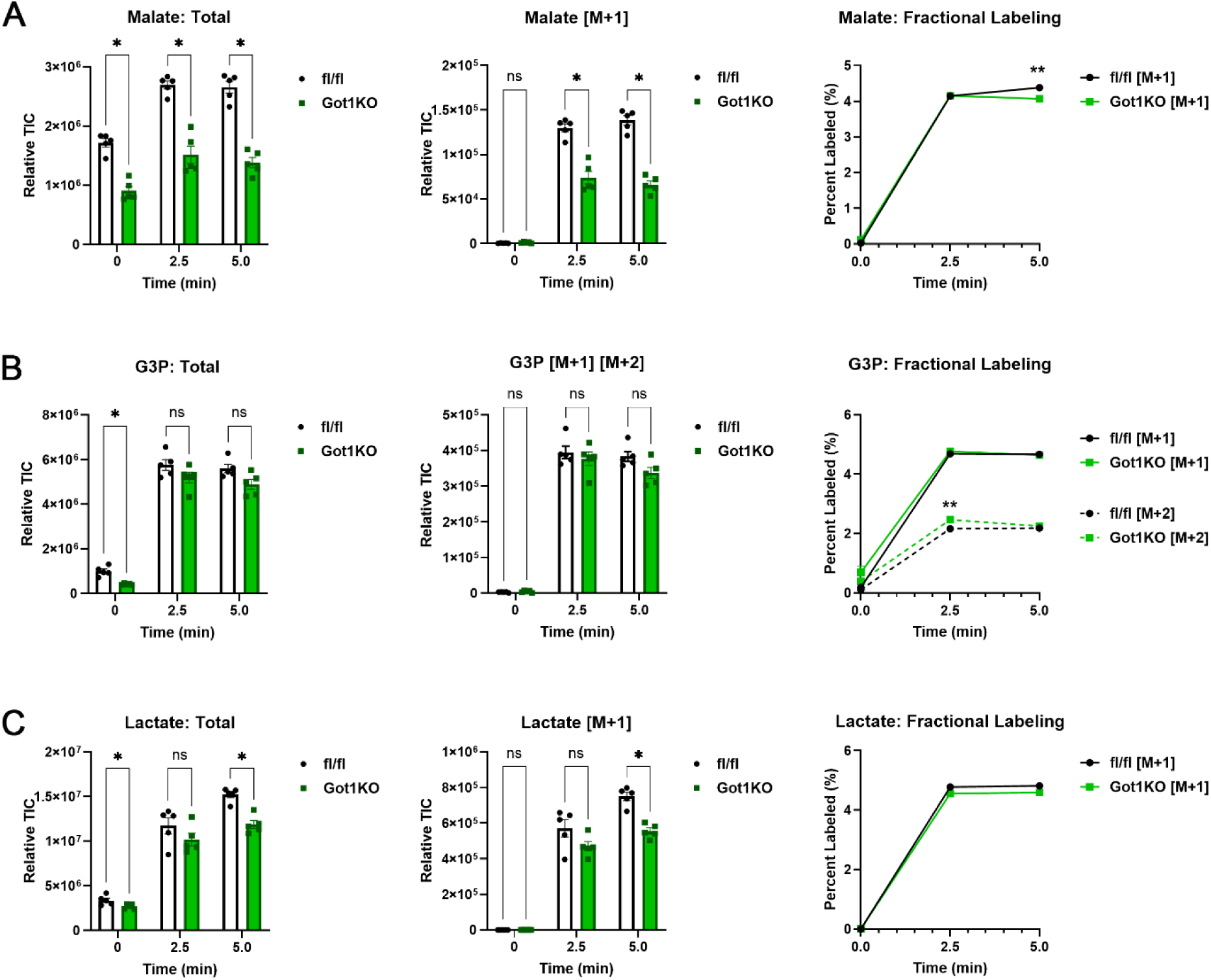
(A) Total and labeled M+1 malate levels and fraction of labeled malate after introducing [4-^2^H] glucose. (B) Total and labeled G3P levels and fraction of labeled M+1 G3P and M+2 G3P after introducing [4-^2^H] glucose. (C) Total and labeled M+1 lactate levels and fraction of labeled lactate after introducing [4-^2^H] glucose. *Got1*fl/fl and *Got1*KO brown adipocytes were treated with isoproterenol for 4h prior to addition of 25 mM [4-^2^H] glucose. The middle panels showing the labeled metabolites are also presented in Figure 7E. All data are presented as the Mean ± SEM. \**p*<0.05, \*\**p*<0.01.

## References

1. Cannon, B., and Nedergaard, J. (2004). Brown adipose tissue: function and physiological significance. Physiol Rev 84, 277–359. 10.1152/physrev.00015.2003 84/1/277 [pii].

2. Nedergaard, J., Golozoubova, V., Matthias, A., Asadi, A., Jacobsson, A., and Cannon, B. (2001). UCP1: the only protein able to mediate adaptive non-shivering thermogenesis and metabolic inefficiency. Biochim Biophys Acta 1504, 82–106. S0005-2728(00)00247-4 [pii].

3. Golozoubova, V., Hohtola, E., Matthias, A., Jacobsson, A., Cannon, B., and Nedergaard, J. (2001). Only UCP1 can mediate adaptive nonshivering thermogenesis in the cold. FASEB J 15, 2048–2050. 10.1096/fj.00-0536fje 00-0536fje [pii].

4. Wu, J., Bostrom, P., Sparks, L.M., Ye, L., Choi, J.H., Giang, A.H., Khandekar, M., Virtanen, K.A., Nuutila, P., Schaart, G., et al. (2012). Beige adipocytes are a distinct type of thermogenic fat cell in mouse and human. Cell 150, 366–376. 10.1016/j.cell.2012.05.016 S0092-8674(12)00595-8 [pii].

5. Himms-Hagen, J., Cui, J., Danforth, E., Jr., Taatjes, D.J., Lang, S.S., Waters, B.L., and Claus, T.H. (1994). Effect of CL-316,243, a thermogenic beta 3-agonist, on energy balance and brown and white adipose tissues in rats. Am J Physiol 266, R1371–1382.

6. Himms-Hagen, J., Melnyk, A., Zingaretti, M.C., Ceresi, E., Barbatelli, G., and Cinti, S. (2000). Multilocular fat cells in WAT of CL-316243-treated rats derive directly from white adipocytes. Am J Physiol Cell Physiol 279, C670–681.

7. Granneman, J.G., Li, P., Zhu, Z., and Lu, Y. (2005). Metabolic and cellular plasticity in white adipose tissue I: effects of beta3-adrenergic receptor activation. Am J Physiol Endocrinol Metab 289, E608–616. 00009.2005 [pii] 10.1152/ajpendo.00009.2005.

8. Cypess, A.M., Lehman, S., Williams, G., Tal, I., Rodman, D., Goldfine, A.B., Kuo, F.C., Palmer, E.L., Tseng, Y.H., Doria, A., et al. (2009). Identification and importance of brown adipose tissue in adult humans. N Engl J Med 360, 1509–1517. 10.1056/NEJMoa0810780 360/15/1509 [pii].

9. Saito, M., Okamatsu-Ogura, Y., Matsushita, M., Watanabe, K., Yoneshiro, T., Nio-Kobayashi, J., Iwanaga, T., Miyagawa, M., Kameya, T., Nakada, K., et al. (2009). High incidence of metabolically active brown adipose tissue in healthy adult humans: effects of cold exposure and adiposity. Diabetes 58, 1526–1531. 10.2337/db09-0530 db09-0530 [pii].

10. van Marken Lichtenbelt, W.D., Vanhommerig, J.W., Smulders, N.M., Drossaerts, J.M., Kemerink, G.J., Bouvy, N.D., Schrauwen, P., and Teule, G.J. (2009). Cold-activated brown adipose tissue in healthy men. N Engl J Med 360, 1500–1508. 10.1056/NEJMoa0808718 360/15/1500 [pii].

11. Chondronikola, M., Volpi, E., Borsheim, E., Porter, C., Annamalai, P., Enerback, S., Lidell, M.E., Saraf, M.K., Labbe, S.M., Hurren, N.M., et al. (2014). Brown adipose tissue improves whole-body glucose homeostasis and insulin sensitivity in humans. Diabetes 63, 4089–4099. 10.2337/db14-0746.

12. Ouellet, V., Labbe, S.M., Blondin, D.P., Phoenix, S., Guerin, B., Haman, F., Turcotte, E.E., Richard, D., and Carpentier, A.C. (2012). Brown adipose tissue oxidative metabolism contributes to energy expenditure during acute cold exposure in humans. J Clin Invest 122, 545–552. 10.1172/JCI60433 60433 [pii].

13. Hanssen, M.J., Hoeks, J., Brans, B., van der Lans, A.A., Schaart, G., van den Driessche, J.J., Jorgensen, J.A., Boekschoten, M.V., Hesselink, M.K., Havekes, B., et al. (2015). Short-term cold acclimation improves insulin sensitivity in patients with type 2 diabetes mellitus. Nat Med 21, 863–865. 10.1038/nm.3891.

14. Cypess, A.M., Weiner, L.S., Roberts-Toler, C., Franquet Elia, E., Kessler, S.H., Kahn, P.A., English, J., Chatman, K., Trauger, S.A., Doria, A., and Kolodny, G.M. (2015). Activation of human brown adipose tissue by a beta3-adrenergic receptor agonist. Cell Metab 21, 33–38. 10.1016/j.cmet.2014.12.009.

15. O’Mara, A.E., Johnson, J.W., Linderman, J.D., Brychta, R.J., McGehee, S., Fletcher, L.A., Fink, Y.A., Kapuria, D., Cassimatis, T.M., Kelsey, N., et al. (2020). Chronic mirabegron treatment increases human brown fat, HDL cholesterol, and insulin sensitivity. J Clin Invest 130, 2209–2219. 10.1172/JCI131126.

16. Townsend, K.L., and Tseng, Y.H. (2014). Brown fat fuel utilization and thermogenesis. Trends Endocrinol Metab 25, 168–177. 10.1016/j.tem.2013.12.004.

17. Fedorenko, A., Lishko, P.V., and Kirichok, Y. (2012). Mechanism of fatty-acid-dependent UCP1 uncoupling in brown fat mitochondria. Cell 151, 400–413. 10.1016/j.cell.2012.09.010.

18. Ma, S.W., and Foster, D.O. (1986). Uptake of glucose and release of fatty acids and glycerol by rat brown adipose tissue in vivo. Can J Physiol Pharmacol 64, 609–614.

19. Isler, D., Hill, H.P., and Meier, M.K. (1987). Glucose metabolism in isolated brown adipocytes under beta-adrenergic stimulation. Quantitative contribution of glucose to total thermogenesis. Biochem J 245, 789–793.

20. Hankir, M.K., and Klingenspor, M. (2018). Brown adipocyte glucose metabolism: a heated subject. EMBO Rep 19. 10.15252/embr.201846404.

21. Jung, S.M., Doxsey, W.G., Le, J., Haley, J.A., Mazuecos, L., Luciano, A.K., Li, H., Jang, C., and Guertin, D.A. (2021). In vivo isotope tracing reveals the versatility of glucose as a brown adipose tissue substrate. Cell Rep 36, 109459. 10.1016/j.celrep.2021.109459.

22. Winther, S., Isidor, M.S., Basse, A.L., Skjoldborg, N., Cheung, A., Quistorff, B., and Hansen, J.B. (2018). Restricting glycolysis impairs brown adipocyte glucose and oxygen consumption. Am J Physiol Endocrinol Metab 314, E214–E223. 10.1152/ajpendo.00218.2017.

23. Bornstein, M.R., Neinast, M.D., Zeng, X., Chu, Q., Axsom, J., Thorsheim, C., Li, K., Blair, M.C., Rabinowitz, J.D., and Arany, Z. (2023). Comprehensive quantification of metabolic flux during acute cold stress in mice. Cell Metab 35, 2077–2092 e2076. 10.1016/j.cmet.2023.09.002.

24. Yu, X.X., Lewin, D.A., Forrest, W., and Adams, S.H. (2002). Cold elicits the simultaneous induction of fatty acid synthesis and beta-oxidation in murine brown adipose tissue: prediction from differential gene expression and confirmation in vivo. FASEB J 16, 155–168. 10.1096/fj.01-0568com 16/2/155 [pii].

25. Verkerke, A.R.P., Wang, D., Yoshida, N., Taxin, Z.H., Shi, X., Zheng, S., Li, Y., Auger, C., Oikawa, S., Yook, J.S., et al. (2024). BCAA-nitrogen flux in brown fat controls metabolic health independent of thermogenesis. Cell 187, 2359–2374 e2318. 10.1016/j.cell.2024.03.030.

26. Yoneshiro, T., Wang, Q., Tajima, K., Matsushita, M., Maki, H., Igarashi, K., Dai, Z., White, P.J., McGarrah, R.W., Ilkayeva, O.R., et al. (2019). BCAA catabolism in brown fat controls energy homeostasis through SLC25A44. Nature 572, 614–619. 10.1038/s41586-019-1503-x.

27. Borst, P. (2020). The malate-aspartate shuttle (Borst cycle): How it started and developed into a major metabolic pathway. IUBMB Life 72, 2241–2259. 10.1002/iub.2367.

28. Pardo, B., Contreras, L., and Satrustegui, J. (2013). De novo Synthesis of Glial Glutamate and Glutamine in Young Mice Requires Aspartate Provided by the Neuronal Mitochondrial Aspartate-Glutamate Carrier Aralar/AGC1. Front Endocrinol (Lausanne) 4, 149. 10.3389/fendo.2013.00149.

29. Lu, M., Zhou, L., Stanley, W.C., Cabrera, M.E., Saidel, G.M., and Yu, X. (2008). Role of the malate-aspartate shuttle on the metabolic response to myocardial ischemia. J Theor Biol 254, 466–475. 10.1016/j.jtbi.2008.05.033.

30. Saheki, T., and Kobayashi, K. (2002). Mitochondrial aspartate glutamate carrier (citrin) deficiency as the cause of adult-onset type II citrullinemia (CTLN2) and idiopathic neonatal hepatitis (NICCD). J Hum Genet 47, 333–341. 10.1007/s100380200046.

31. Rubi, B., del Arco, A., Bartley, C., Satrustegui, J., and Maechler, P. (2004). The malate-aspartate NADH shuttle member Aralar1 determines glucose metabolic fate, mitochondrial activity, and insulin secretion in beta cells. J Biol Chem 279, 55659–55666. 10.1074/jbc.M409303200.

32. Schantz, P.G., Sjoberg, B., and Svedenhag, J. (1986). Malate-aspartate and alpha-glycerophosphate shuttle enzyme levels in human skeletal muscle: methodological considerations and effect of endurance training. Acta Physiol Scand 128, 397–407. 10.1111/j.1748-1716.1986.tb07993.x.

33. Chang, J.S., Ghosh, S., Newman, S., and Salbaum, J.M. (2018). A map of the PGC-1alpha- and NT-PGC-1alpha-regulated transcriptional network in brown adipose tissue. Sci Rep 8, 7876. 10.1038/s41598-018-26244-4.

34. Perdikari, A., Leparc, G.G., Balaz, M., Pires, N.D., Lidell, M.E., Sun, W., Fernandez-Albert, F., Muller, S., Akchiche, N., Dong, H., et al. (2018). BATLAS: Deconvoluting Brown Adipose Tissue. Cell Rep 25, 784–797 e784. 10.1016/j.celrep.2018.09.044.

35. Zhang, Y., Huypens, P., Adamson, A.W., Chang, J.S., Henagan, T.M., Lenard, N.R., Burk, D., Klein, J., Perwitz, N., Shin, J., et al. (2009). Alternative mRNA splicing produces a novel biologically active short isoform of PGC-1{alpha}. J Biol Chem 284, 32813–32826.

36. Chang, J.S., Fernand, V., Zhang, Y., Shin, J., Jun, H.J., Joshi, Y., and Gettys, T.W. (2012). NT-PGC-1alpha protein is sufficient to link beta3-adrenergic receptor activation to transcriptional and physiological components of adaptive thermogenesis. J Biol Chem 287, 9100–9111. M111.320200 [pii] 10.1074/jbc.M111.320200.

37. Dufour, C.R., Wilson, B.J., Huss, J.M., Kelly, D.P., Alaynick, W.A., Downes, M., Evans, R.M., Blanchette, M., and Giguere, V. (2007). Genome-wide orchestration of cardiac functions by the orphan nuclear receptors ERRalpha and gamma. Cell Metab 5, 345–356. 10.1016/j.cmet.2007.03.007.

38. Schreiber, S.N., Emter, R., Hock, M.B., Knutti, D., Cardenas, J., Podvinec, M., Oakeley, E.J., and Kralli, A. (2004). The estrogen-related receptor alpha (ERRalpha) functions in PPARgamma coactivator 1alpha (PGC-1alpha)-induced mitochondrial biogenesis. Proc Natl Acad Sci U S A 101, 6472–6477.

39. Schreiber, S.N., Knutti, D., Brogli, K., Uhlmann, T., and Kralli, A. (2003). The transcriptional coactivator PGC-1 regulates the expression and activity of the orphan nuclear receptor estrogen-related receptor alpha (ERRalpha). J Biol Chem 278, 9013–9018. 10.1074/jbc.M212923200.

40. Uldry, M., Yang, W., St-Pierre, J., Lin, J., Seale, P., and Spiegelman, B.M. (2006). Complementary action of the PGC-1 coactivators in mitochondrial biogenesis and brown fat differentiation. Cell Metab 3, 333–341.

41. Jun, H.J., Joshi, Y., Patil, Y., Noland, R.C., and Chang, J.S. (2014). NT-PGC-1alpha activation attenuates high-fat diet-induced obesity by enhancing brown fat thermogenesis and adipose tissue oxidative metabolism. Diabetes 63, 3615–3625. 10.2337/db13-1837 db13-1837 [pii].

42. Kim, J., Park, M.S., Ha, K., Park, C., Lee, J., Mynatt, R.L., and Chang, J.S. (2018). NT-PGC-1alpha deficiency decreases mitochondrial FA oxidation in brown adipose tissue and alters substrate utilization in vivo. J Lipid Res 59, 1660–1670. 10.1194/jlr.M085647.

43. Kong, X., Banks, A., Liu, T., Kazak, L., Rao, R.R., Cohen, P., Wang, X., Yu, S., Lo, J.C., Tseng, Y.H., et al. (2014). IRF4 is a key thermogenic transcriptional partner of PGC-1alpha. Cell 158, 69–83. 10.1016/j.cell.2014.04.049 S0092-8674(14)00723-5 [pii].

44. Shibata, H., Perusse, F., Vallerand, A., and Bukowiecki, L.J. (1989). Cold exposure reverses inhibitory effects of fasting on peripheral glucose uptake in rats. Am J Physiol 257, R96–101. 10.1152/ajpregu.1989.257.1.R96.

45. Albert, V., Svensson, K., Shimobayashi, M., Colombi, M., Munoz, S., Jimenez, V., Handschin, C., Bosch, F., and Hall, M.N. (2016). mTORC2 sustains thermogenesis via Akt-induced glucose uptake and glycolysis in brown adipose tissue. EMBO Mol Med 8, 232–246. 10.15252/emmm.201505610 emmm.201505610 [pii].

46. Bartelt, A., Bruns, O.T., Reimer, R., Hohenberg, H., Ittrich, H., Peldschus, K., Kaul, M.G., Tromsdorf, U.I., Weller, H., Waurisch, C., et al. (2011). Brown adipose tissue activity controls triglyceride clearance. Nat Med 17, 200–205. 10.1038/nm.2297 nm.2297 [pii].

47. Gasparetti, A.L., de Souza, C.T., Pereira-da-Silva, M., Oliveira, R.L., Saad, M.J., Carneiro, E.M., and Velloso, L.A. (2003). Cold exposure induces tissue-specific modulation of the insulin-signalling pathway in Rattus norvegicus. J Physiol 552, 149–162. 10.1113/jphysiol.2003.050369.

48. Wang, L., Jin, Q., Lee, J.E., Su, I.H., and Ge, K. (2010). Histone H3K27 methyltransferase Ezh2 represses Wnt genes to facilitate adipogenesis. Proc Natl Acad Sci U S A 107, 7317–7322. 10.1073/pnas.1000031107.

49. Eto, K., Tsubamoto, Y., Terauchi, Y., Sugiyama, T., Kishimoto, T., Takahashi, N., Yamauchi, N., Kubota, N., Murayama, S., Aizawa, T., et al. (1999). Role of NADH shuttle system in glucose-induced activation of mitochondrial metabolism and insulin secretion. Science 283, 981–985.

50. Hung, Y.P., Albeck, J.G., Tantama, M., and Yellen, G. (2011). Imaging cytosolic NADH-NAD(+) redox state with a genetically encoded fluorescent biosensor. Cell Metab 14, 545–554. 10.1016/j.cmet.2011.08.012.

51. Hung, Y.P., and Yellen, G. (2014). Live-cell imaging of cytosolic NADH-NAD+ redox state using a genetically encoded fluorescent biosensor. Methods Mol Biol 1071, 83–95. 10.1007/978-1-62703-622-1_7.

52. Diaz-Garcia, C.M., Mongeon, R., Lahmann, C., Koveal, D., Zucker, H., and Yellen, G. (2017). Neuronal Stimulation Triggers Neuronal Glycolysis and Not Lactate Uptake. Cell Metab 26, 361–374 e364. 10.1016/j.cmet.2017.06.021.

53. Hung, Y.P., Teragawa, C., Kosaisawe, N., Gillies, T.E., Pargett, M., Minguet, M., Distor, K., Rocha-Gregg, B.L., Coloff, J.L., Keibler, M.A., et al. (2017). Akt regulation of glycolysis mediates bioenergetic stability in epithelial cells. Elife 6. 10.7554/eLife.27293.

54. Hanse, E.A., Ruan, C., Kachman, M., Wang, D., Lowman, X.H., and Kelekar, A. (2017). Cytosolic malate dehydrogenase activity helps support glycolysis in actively proliferating cells and cancer. Oncogene 36, 3915–3924. 10.1038/onc.2017.36.

55. Liu, L., Shah, S., Fan, J., Park, J.O., Wellen, K.E., and Rabinowitz, J.D. (2016). Malic enzyme tracers reveal hypoxia-induced switch in adipocyte NADPH pathway usage. Nat Chem Biol 12, 345–352. 10.1038/nchembio.2047.

56. Lewis, C.A., Parker, S.J., Fiske, B.P., McCloskey, D., Gui, D.Y., Green, C.R., Vokes, N.I., Feist, A.M., Vander Heiden, M.G., and Metallo, C.M. (2014). Tracing compartmentalized NADPH metabolism in the cytosol and mitochondria of mammalian cells. Mol Cell 55, 253–263. 10.1016/j.molcel.2014.05.008.

57. Wu, M., Neilson, A., Swift, A.L., Moran, R., Tamagnine, J., Parslow, D., Armistead, S., Lemire, K., Orrell, J., Teich, J., et al. (2007). Multiparameter metabolic analysis reveals a close link between attenuated mitochondrial bioenergetic function and enhanced glycolysis dependency in human tumor cells. Am J Physiol Cell Physiol 292, C125–136. 10.1152/ajpcell.00247.2006.

58. Pelletier, A., Joly, E., Prentki, M., and Coderre, L. (2005). Adenosine 5’-monophosphate-activated protein kinase and p38 mitogen-activated protein kinase participate in the stimulation of glucose uptake by dinitrophenol in adult cardiomyocytes. Endocrinology 146, 2285–2294. 10.1210/en.2004-1565.

59. Tao, H., Zhang, Y., Zeng, X., Shulman, G.I., and Jin, S. (2014). Niclosamide ethanolamine-induced mild mitochondrial uncoupling improves diabetic symptoms in mice. Nat Med 20, 1263–1269. 10.1038/nm.3699.

60. Axelrod, C.L., King, W.T., Davuluri, G., Noland, R.C., Hall, J., Hull, M., Dantas, W.S., Zunica, E.R., Alexopoulos, S.J., Hoehn, K.L., et al. (2020). BAM15-mediated mitochondrial uncoupling protects against obesity and improves glycemic control. EMBO Mol Med 12, e12088. 10.15252/emmm.202012088.

61. Pulinilkunnil, T., He, H., Kong, D., Asakura, K., Peroni, O.D., Lee, A., and Kahn, B.B. (2011). Adrenergic regulation of AMP-activated protein kinase in brown adipose tissue in vivo. J Biol Chem 286, 8798–8809. 10.1074/jbc.M111.218719.

62. Gauthier, M.S., Miyoshi, H., Souza, S.C., Cacicedo, J.M., Saha, A.K., Greenberg, A.S., and Ruderman, N.B. (2008). AMP-activated protein kinase is activated as a consequence of lipolysis in the adipocyte: potential mechanism and physiological relevance. J Biol Chem 283, 16514–16524. 10.1074/jbc.M708177200.

63. Mihaylova, M.M., and Shaw, R.J. (2011). The AMPK signalling pathway coordinates cell growth, autophagy and metabolism. Nat Cell Biol 13, 1016–1023. 10.1038/ncb2329.

64. Inokuma, K., Ogura-Okamatsu, Y., Toda, C., Kimura, K., Yamashita, H., and Saito, M. (2005). Uncoupling protein 1 is necessary for norepinephrine-induced glucose utilization in brown adipose tissue. Diabetes 54, 1385–1391. 10.2337/diabetes.54.5.1385.

65. Mottillo, E.P., Desjardins, E.M., Crane, J.D., Smith, B.K., Green, A.E., Ducommun, S., Henriksen, T.I., Rebalka, I.A., Razi, A., Sakamoto, K., et al. (2016). Lack of Adipocyte AMPK Exacerbates Insulin Resistance and Hepatic Steatosis through Brown and Beige Adipose Tissue Function. Cell Metab 24, 118–129. 10.1016/j.cmet.2016.06.006.

66. Chen, Z.P., McConell, G.K., Michell, B.J., Snow, R.J., Canny, B.J., and Kemp, B.E. (2000). AMPK signaling in contracting human skeletal muscle: acetyl-CoA carboxylase and NO synthase phosphorylation. Am J Physiol Endocrinol Metab 279, E1202–1206. 10.1152/ajpendo.2000.279.5.E1202.

67. Abbud, W., Habinowski, S., Zhang, J.Z., Kendrew, J., Elkairi, F.S., Kemp, B.E., Witters, L.A., and Ismail-Beigi, F. (2000). Stimulation of AMP-activated protein kinase (AMPK) is associated with enhancement of Glut1-mediated glucose transport. Arch Biochem Biophys 380, 347–352. 10.1006/abbi.2000.1935.

68. McGee, S.L., van Denderen, B.J., Howlett, K.F., Mollica, J., Schertzer, J.D., Kemp, B.E., and Hargreaves, M. (2008). AMP-activated protein kinase regulates GLUT4 transcription by phosphorylating histone deacetylase 5. Diabetes 57, 860–867. 10.2337/db07-0843.

69. Kurth-Kraczek, E.J., Hirshman, M.F., Goodyear, L.J., and Winder, W.W. (1999). 5’ AMP-activated protein kinase activation causes GLUT4 translocation in skeletal muscle. Diabetes 48, 1667–1671. 10.2337/diabetes.48.8.1667.

70. Fryer, L.G., Foufelle, F., Barnes, K., Baldwin, S.A., Woods, A., and Carling, D. (2002). Characterization of the role of the AMP-activated protein kinase in the stimulation of glucose transport in skeletal muscle cells. Biochem J 363, 167–174. 10.1042/0264-6021:3630167.

71. Marsin, A.S., Bertrand, L., Rider, M.H., Deprez, J., Beauloye, C., Vincent, M.F., Van den Berghe, G., Carling, D., and Hue, L. (2000). Phosphorylation and activation of heart PFK-2 by AMPK has a role in the stimulation of glycolysis during ischaemia. Curr Biol 10, 1247–1255. 10.1016/s0960-9822(00)00742-9.

72. Altshuler-Keylin, S., Shinoda, K., Hasegawa, Y., Ikeda, K., Hong, H., Kang, Q., Yang, Y., Perera, R.M., Debnath, J., and Kajimura, S. (2016). Beige Adipocyte Maintenance Is Regulated by Autophagy-Induced Mitochondrial Clearance. Cell Metab 24, 402–419. 10.1016/j.cmet.2016.08.002.

73. Sambeat, A., Gulyaeva, O., Dempersmier, J., Tharp, K.M., Stahl, A., Paul, S.M., and Sul, H.S. (2016). LSD1 Interacts with Zfp516 to Promote UCP1 Transcription and Brown Fat Program. Cell Rep 15, 2536–2549. 10.1016/j.celrep.2016.05.019.

74. Matesanz, N., Nikolic, I., Leiva, M., Pulgarin-Alfaro, M., Santamans, A.M., Bernardo, E., Mora, A., Herrera-Melle, L., Rodriguez, E., Beiroa, D., et al. (2018). p38alpha blocks brown adipose tissue thermogenesis through p38delta inhibition. PLoS Biol 16, e2004455. 10.1371/journal.pbio.2004455.

75. Ohashi, N., Morino, K., Ida, S., Sekine, O., Lemecha, M., Kume, S., Park, S.Y., Choi, C.S., Ugi, S., and Maegawa, H. (2017). Pivotal Role of O-GlcNAc Modification in Cold-Induced Thermogenesis by Brown Adipose Tissue Through Mitochondrial Biogenesis. Diabetes 66, 2351–2362. 10.2337/db16-1427.

76. Bricker, D.K., Taylor, E.B., Schell, J.C., Orsak, T., Boutron, A., Chen, Y.C., Cox, J.E., Cardon, C.M., Van Vranken, J.G., Dephoure, N., et al. (2012). A mitochondrial pyruvate carrier required for pyruvate uptake in yeast, Drosophila, and humans. Science 337, 96–100. 10.1126/science.1218099.

77. Herzig, S., Raemy, E., Montessuit, S., Veuthey, J.L., Zamboni, N., Westermann, B., Kunji, E.R., and Martinou, J.C. (2012). Identification and functional expression of the mitochondrial pyruvate carrier. Science 337, 93–96. 10.1126/science.1218530.

78. Gerhart-Hines, Z., Rodgers, J.T., Bare, O., Lerin, C., Kim, S.H., Mostoslavsky, R., Alt, F.W., Wu, Z., and Puigserver, P. (2007). Metabolic control of muscle mitochondrial function and fatty acid oxidation through SIRT1/PGC-1alpha. EMBO J 26, 1913–1923. 7601633 [pii] 10.1038/sj.emboj.7601633.

79. Lagouge, M., Argmann, C., Gerhart-Hines, Z., Meziane, H., Lerin, C., Daussin, F., Messadeq, N., Milne, J., Lambert, P., Elliott, P., et al. (2006). Resveratrol improves mitochondrial function and protects against metabolic disease by activating SIRT1 and PGC-1alpha. Cell 127, 1109–1122. 10.1016/j.cell.2006.11.013.

80. Nemoto, S., Fergusson, M.M., and Finkel, T. (2005). SIRT1 functionally interacts with the metabolic regulator and transcriptional coactivator PGC-1{alpha}. J Biol Chem 280, 16456–16460. M501485200 [pii] 10.1074/jbc.M501485200.

81. Adina-Zada, A., Zeczycki, T.N., St Maurice, M., Jitrapakdee, S., Cleland, W.W., and Attwood, P.V. (2012). Allosteric regulation of the biotin-dependent enzyme pyruvate carboxylase by acetyl-CoA. Biochem Soc Trans 40, 567–572. 10.1042/BST20120041.

82. Houstek, J., Cannon, B., and Lindberg, O. (1975). Gylcerol-3-phosphate shuttle and its function in intermediary metabolism of hamster brown-adipose tissue. Eur J Biochem 54, 11–18.

83. Nguyen, H.P., Yi, D., Lin, F., Viscarra, J.A., Tabuchi, C., Ngo, K., Shin, G., Lee, A.Y., Wang, Y., and Sul, H.S. (2020). Aifm2, a NADH Oxidase, Supports Robust Glycolysis and Is Required for Cold-and Diet-Induced Thermogenesis. Mol Cell 77, 600–617 e604. 10.1016/j.molcel.2019.12.002.

84. Moura, M.A., Festuccia, W.T., Kawashita, N.H., Garofalo, M.A., Brito, S.R., Kettelhut, I.C., and Migliorini, R.H. (2005). Brown adipose tissue glyceroneogenesis is activated in rats exposed to cold. Pflugers Arch 449, 463–469. 10.1007/s00424-004-1353-7.

85. Claflin, K.E., Flippo, K.H., Sullivan, A.I., Naber, M.C., Zhou, B., Neff, T.J., Jensen-Cody, S.O., and Potthoff, M.J. (2022). Conditional gene targeting using UCP1-Cre mice directly targets the central nervous system beyond thermogenic adipose tissues. Mol Metab 55, 101405. 10.1016/j.molmet.2021.101405.

86. Chang, J.S., Huypens, P., Zhang, Y., Black, C., Kralli, A., and Gettys, T.W. (2010). Regulation of NT-PGC-1alpha subcellular localization and function by protein kinase A-dependent modulation of nuclear export by CRM1. J Biol Chem 285, 18039–18050. M109.083121 [pii] 10.1074/jbc.M109.083121.

87. Agrawal, S., Kumar, S., Sehgal, R., George, S., Gupta, R., Poddar, S., Jha, A., and Pathak, S. (2019). El-MAVEN: A Fast, Robust, and User-Friendly Mass Spectrometry Data Processing Engine for Metabolomics. Methods Mol Biol 1978, 301–321. 10.1007/978-1-4939-9236-2_19.

88. Heinrich, P., Kohler, C., Ellmann, L., Kuerner, P., Spang, R., Oefner, P.J., and Dettmer, K. (2018). Correcting for natural isotope abundance and tracer impurity in MS-, MS/MS- and high-resolution-multiple-tracer-data from stable isotope labeling experiments with IsoCorrectoR. Sci Rep 8, 17910. 10.1038/s41598-018-36293-4.

89. Huynh, F.K., Green, M.F., Koves, T.R., and Hirschey, M.D. (2014). Measurement of fatty acid oxidation rates in animal tissues and cell lines. Methods Enzymol 542, 391–405. 10.1016/B978-0-12-416618-9.00020-0.

90. Gray, L.R., Rauckhorst, A.J., and Taylor, E.B. (2016). A Method for Multiplexed Measurement of Mitochondrial Pyruvate Carrier Activity. J Biol Chem 291, 7409–7417. 10.1074/jbc.M115.711663.

91. Mookerjee, S.A., and Brand, M.D. (2015). Measurement and Analysis of Extracellular Acid Production to Determine Glycolytic Rate. J Vis Exp, e53464. 10.3791/53464.

92. Commins, S.P., Watson, P.M., Padgett, M.A., Dudley, A., Argyropoulos, G., and Gettys, T.W. (1999). Induction of uncoupling protein expression in brown and white adipose tissue by leptin. Endocrinology 140, 292–300. 10.1210/endo.140.1.6399.

93. Wang, X., and Seed, B. (2003). A PCR primer bank for quantitative gene expression analysis. Nucleic Acids Res 31, e154. 10.1093/nar/gng154.

